# Transcription Factor Dynamics in Cross-Regulation of Plant Hormone Signaling Pathways

**DOI:** 10.1101/2023.03.07.531630

**Authors:** Lingling Yin, Mark Zander, Shao-shan Carol Huang, Mingtang Xie, Liang Song, J. Paola Saldierna Guzmán, Elizabeth Hann, Bhuvana K. Shanbhag, Sophia Ng, Siddhartha Jain, Bart J. Janssen, Natalie M. Clark, Justin W. Walley, Travis Beddoe, Ziv Bar-Joseph, Mathew G. Lewsey, Joseph R. Ecker

## Abstract

Cross-regulation between hormone signaling pathways is indispensable for plant growth and development. However, the molecular mechanisms by which multiple hormones interact and co-ordinate activity need to be understood. Here, we generated a cross-regulation network explaining how hormone signals are integrated from multiple pathways in etiolated Arabidopsis (*Arabidopsis thaliana*) seedlings. To do so we comprehensively characterized transcription factor activity during plant hormone responses and reconstructed dynamic transcriptional regulatory models for six hormones; abscisic acid, brassinosteroid, ethylene, jasmonic acid, salicylic acid and strigolactone/karrikin. These models incorporated target data for hundreds of transcription factors and thousands of protein-protein interactions. Each hormone recruited different combinations of transcription factors, a subset of which were shared between hormones. Hub target genes existed within hormone transcriptional networks, exhibiting transcription factor activity themselves. In addition, a group of MITOGEN-ACTIVATED PROTEIN KINASES (MPKs) were identified as potential key points of cross-regulation between multiple hormones. Accordingly, the loss of function of one of these (MPK6) disrupted the global proteome, phosphoproteome and transcriptome during hormone responses. Lastly, we determined that all hormones drive substantial alternative splicing that has distinct effects on the transcriptome compared with differential gene expression, acting in early hormone responses. These results provide a comprehensive understanding of the common features of plant transcriptional regulatory pathways and how cross-regulation between hormones acts upon gene expression.

## Introduction

Cross-regulation between hormone signaling pathways is fundamental to plant growth and development. It allows plants to monitor a multitude of external environmental and internal cellular signals, process and integrate this information, then initiate appropriate responses. This enables plants to exhibit plastic development, adapt to their local environment, optimize resource usage and respond to stresses (Jaillais and Chory, 2010; Vanstraelen and Benková, 2012; Aerts et al., 2020; Khan et al., 2020). The growth-defense trade-off is a well-known example, whereby plants experiencing pathogen attack prioritize resource allocation to defense at the expense of growth (Karasov et al., 2017; Guo et al., 2018; Figueroa-Macías et al., 2021). However, this trade-off can be condition-dependent, with plants growing in nutrient-rich conditions not necessarily needing to prioritize one response over the other (Figueroa-Macías et al., 2021).

Each plant hormone has a recognized, distinct signaling pathway (Huang et al., 2017; Binder, 2020; Bürger and Chory, 2020; Chen et al., 2020; Ding and Ding, 2020; Nolan et al., 2020; Yao and Waters, 2020). Cross-regulation between these pathways occurs during signal transduction and regulation of transcription (Jaillais and Chory, 2010). Cross-regulation of transcription can occur through transcription factors (TFs) shared between pathways and by regulation of shared target genes by independent TFs. The latter may be considered less common because a minority of genes is shared between the transcriptional responses to different hormones (Nemhauser et al., 2006). The DELLA and JASMONATE-ZIM DOMAIN (JAZ) proteins and NONEXPRESSER OF PR GENES 1 (NPR1) are classic examples of hormone cross-regulation. In each case, these proteins are primarily regulated by one hormone and are involved in the regulation of that hormone’s pathway, but they also influence other hormone signaling pathways (Achard et al., 2003; Fu and Harberd, 2003; Hou et al., 2010; Yang et al., 2012). Recent research demonstrates that there are multiple points of contact between most plant signaling pathways, indicating hormone signaling pathways likely operate as a highly connected network that permits complex exchange and processing of information (Altmann et al., 2020).

The expression of thousands of genes changes in response to a hormone stimulus (Nemhauser et al., 2006). These expression changes are dynamic over time, with great diversity between the expression patterns of genes (Chang et al., 2013; Song et al., 2016; Xie et al., 2018; Zander et al., 2020). Different TFs act at different times during responses to regulate genes in this dynamic manner. Expression of tens to hundreds of TFs is regulated by the hormones abscisic acid (ABA), ethylene (ET) and jasmonic acid (JA) and it is likely all hormones do similarly (Chang et al., 2013; Song et al., 2016; Zander et al., 2020; Clark et al., 2021). Individual TFs often target hundreds to thousands of genes and individual genes may be targeted by multiple TFs. This enables dynamic and complex expression patterns but presents a substantial problem in determining which TFs regulate these patterns (Chang et al., 2013; Song et al., 2016).

The extent of alternative splicing in hormone responses is not fully understood (Zander et al., 2020). Alternative splicing and variant isoform usage diversify the proteome by permitting individual genes to encode multiple proteins that may vary in structure and function (Syed et al., 2012; Filichkin et al., 2015; Hartmann et al., 2016; Calixto et al., 2018). For example, variant isoforms of the JAZ repressor JAZ10, one encoding an active form of the protein and one a dominant negative form, have an important role in regulating the core JA signaling pathway (Yan et al., 2007; Chung et al., 2009; Moreno et al., 2013). More recently, greater than 100 genes were determined to switch dominant isoforms during a JA response in etiolated Arabidopsis seedlings (Zander et al., 2020). However, whether or not variant isoform usage is a core feature of hormone signaling pathways is unknown.

In this study we set out to understand how cross-regulation of dynamic transcriptional responses to hormones occurs in the etiolated Arabidopsis seedling, a well-characterized model for plant hormonal signaling and development. We did so by analyzing transcriptome dynamics following stimulation of the ABA, brassinosteroid (BR), ET, JA, salicylic acid (SA) and strigolactone/karrikin (SL/KAR) signaling pathways. We identified genes that undergo differential alternative splicing during hormone responses, which extends our understanding of hormone signaling complexity. We also determined the *in vivo* target genes of key TFs then developed models of hormone transcriptional responses that integrated target data for hundreds of other TFs and thousands of protein-protein interactions. The multi-hormone transcriptional model we have developed helps explain how hormones integrate signals from multiple pathways to dynamically cross-regulate gene expression.

## Results

### Reconstruction of dynamic hormone transcriptional regulatory pathways

The major aims of our study were to determine the extent to which cross-regulation of transcription occurs between hormone signaling pathways and to identify components responsible for this cross-regulation. We first reconstructed the dynamic hormone transcriptional regulatory pathways for ABA, BR, ET, JA, SA and SL/KAR from hormone receptors, through signal transduction to TF-gene binding and differential gene expression (Fig. 1).

**Figure 1.**
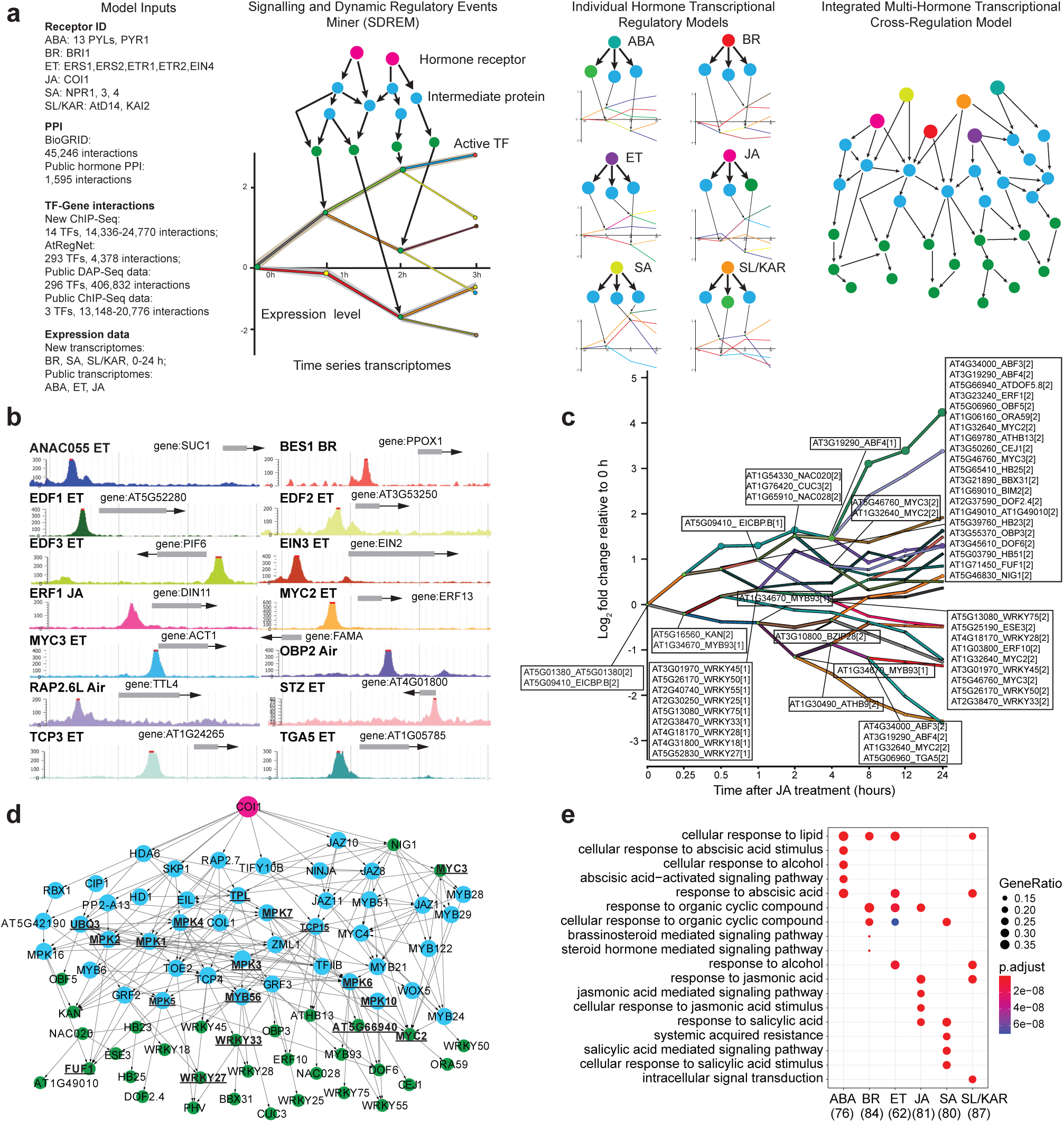
Overview of hormone transcriptional regulatory models reconstructed using the SDREM modeling framework. **a**, The modeling approach underlying our hormone cross-regulation network. Model inputs lists data generated by our lab or from published studies used in the models. SDREM integrates TF-gene interactions and PPIs with time series expression data to build models in an iterative manner. It first identifies active TFs that bind cohorts of co-regulated genes, then searches for paths from hormone receptor(s) to these TFs. Individual models were generated for each hormone of ABA, BR, ET, JA, SA and SL/KAR. These were combined to give the integrated model. **b**, Genome browser screen shot visualizing representative target genes from ChIP-seq samples of 14 TFs. **c**, The regulatory network of the JA model. The network displays all predicted active TFs at each branch point (node) and the bars indicate co-expressed and co-regulated genes. [1] indicates the TF primarily controls the lower path out of the split and [2] is for the higher path. The y-axis is the log2 fold change in expression relative to expression at 0 h. **d**, The JA signaling pathway reconstructed by SDREM. The JA receptor, intermediate proteins and active TFs are indicated by magenta, blue and green nodes respectively. The proteins shared by at least 4 hormone pathways, are in black bold text and have underlined names. **e**, Top five significantly enriched (p.adjust < 0.05) gene ontology biological process terms amongst the predicted nodes of the reconstructed signaling pathway for each hormone.

We generated a model of each hormone which described regulation of the transcriptome over time following treatment with that hormone (Fig. 1a). This was achieved by analyzing time-series transcriptomes from our own newly generated data and published data (BR, ET, JA, SA, SL/KAR, 0-24 h after treatment; ABA, 0-60 h; data sources detailed in Methods) (Extended Data Fig. 1, 2; Supplementary Table 1). Next, we applied Signaling and Dynamic Regulatory Events Miner (SDREM) modeling to reconstruct individual hormone pathways (Fig. 1a) (Gitter and Bar-Joseph, 2013; Gitter et al., 2013; Gitter and Bar-Joseph, 2016). SDREM first identifies dynamic regulation by searching for TFs that bind groups of co-regulated genes during the hormone transcriptional response. Next, it searches for paths from the receptor(s) of that hormone through protein-protein interactions to these regulating TFs, inferring that the paths are mechanisms that may activate the TF during the hormone response. SDREM then iteratively refines the network by penalizing TFs that are not supported by signaling pathways from the receptors.

SDREM modeling requires extensive data about TF-target gene interactions and protein-protein interactions to reconstruct signaling pathways. To enable this, we determined the in vivo target genes of 14 hormone TFs by chromatin immunoprecipitation sequencing (ChIP-seq) to use in model construction, selected because they had substantial pre-existing evidence supporting their key roles in hormone signaling (Fig. 1b; Supplementary Table 2, 3). These were combined with public TF-target gene data for a further 516 TFs (Yilmaz et al., 2010; Song et al., 2016; Narsai et al., 2017; Zander et al., 2020). The known receptors for each hormone and extensive public Arabidopsis protein-protein interaction data (46,841 interactions) were used to build signaling pathways (Stark et al., 2006). We successfully reconstructed the transcriptional regulatory pathways for all six hormones (ABA, BR, ET, JA, SA, SL/KAR) using this approach, demonstrated by each model being enriched for known components of the relevant hormone signaling pathway (Fig. 1c, d, e; Extended Data Fig. 3, 4, 5, 6; Supplementary Table 4, 5). The SDREM models illustrate that each hormone remodels the transcriptome rapidly - within 15 minutes (BR, ET, JA, SA, SL/KAR) or 1 hour (ABA) - of perception of that hormone. Furthermore, transcriptome remodeling is extensive and dynamic, affecting thousands of genes over 24 h (Extended Data Fig. 2).

### The populations of TFs employed by each hormone differ but share some components

We examined the extent to which individual hormones used different TFs to control gene expression. We did so by first identifying shared and unique predicted regulatory TFs within each hormone model. A minority of unique TFs was present in the models of every hormone (19.5% to 34.7% of TFs per model, Fig. 2a; Supplementary Table 6). This is consistent with the prior observation that most genes differentially expressed in response to individual hormones are not shared between different hormones (Nemhauser et al., 2006) (Supplementary Table 7). However, there were also TFs shared between multiple hormone models (Fig. 2a). Fourteen TFs were shared between the models of 4 or more hormones. For example, MYC2, MYC3 and WRKY33 were all predicted regulators in the BR, JA, SA and SL/KAR models (Fig. 2a). Overall, we observed that each hormone used distinct combinations of TF families to regulate transcription. This was demonstrated by the relative enrichment of TF families between hormone models (Fig. 2b; Supplementary Table 6). These results indicate that although the responses to ABA, BR, ET, JA, SA and SL/KAR share some TFs, their unique transcriptional regulatory pathways are established by recruiting different combinations of TFs.

**Figure 2.**
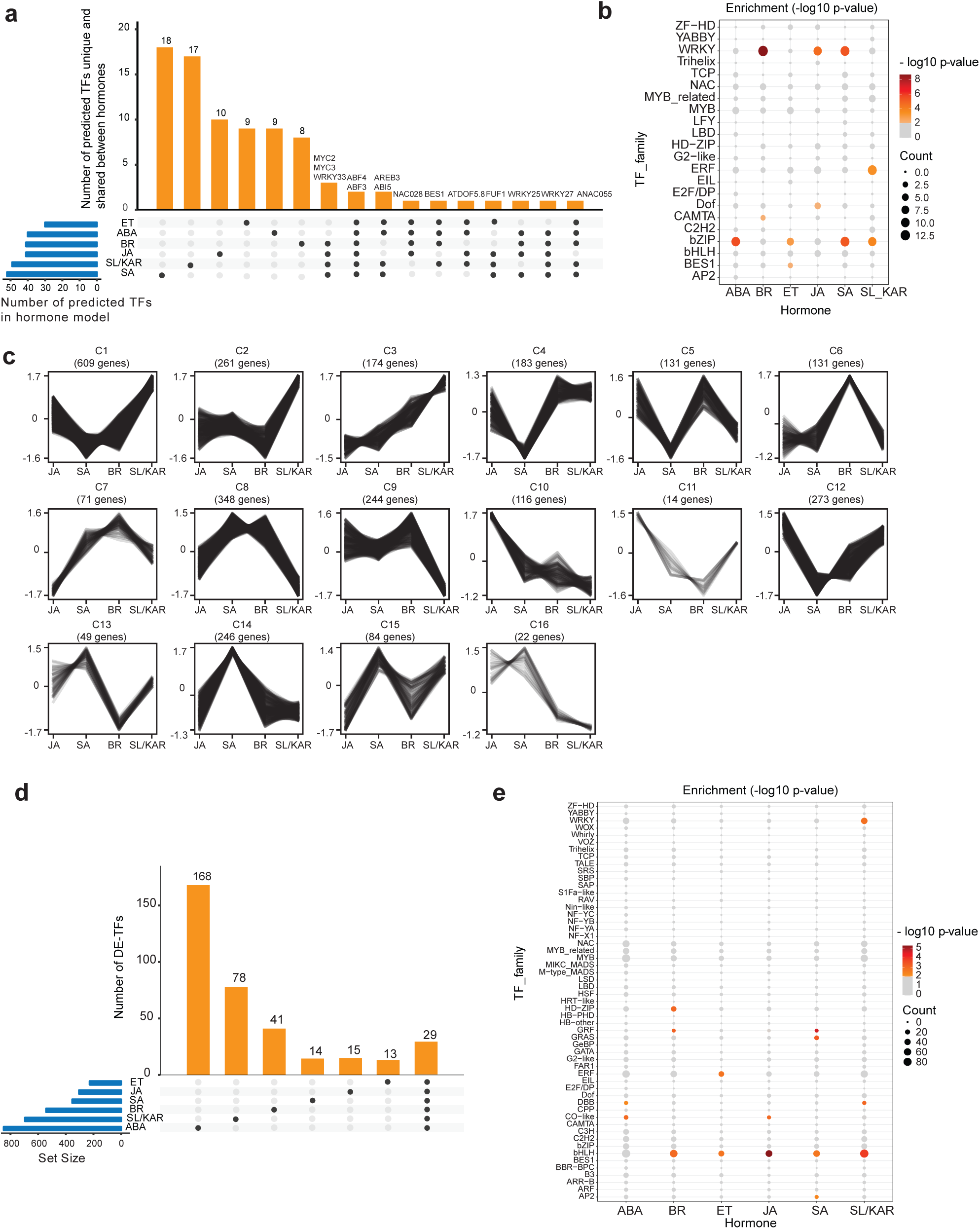
Different TFs regulate the response to each hormone. **a**, The number of active TFs unique to and shared between all six hormone models. Names of TFs shared between 4 or more hormone models are labelled at the top of respective columns. **b**, Significantly enriched TF families found within each hormone model (p-value < 0.01; hypergeometric test). The size and colour of each circle represents per-family TF count and enrichment p-value range respectively. **c**, K-means clustering of expression of MYC2 target genes during JA, SA, BR and SL/KAR hormone responses. Expression is given as normalized transcripts per million (TPM). **d**, The number of unique and shared differentially expressed TFs (DE-TFs) between six hormones. **e**, Significantly enriched TF families amongst DE-TFs for each hormone (p-value < 0.01; hypergeometric test). The size and colour of each circle represents per-family TF count and enrichment p-value range respectively.

TFs shared between multiple hormones might perform the same function for each hormone, meaning that they regulate genes in the same manner in all conditions. However, we observed that shared TFs were predicted to regulate gene expression at different times post-treatment in each hormone response and were associated with up and down-regulated genes (Extended Data Fig. 7a). This suggested the former proposal was unlikely. Alternatively, shared TFs might regulate a common set of genes in different manners - promoting expression for one hormone, repressing expression for another - or regulate distinct sets of genes for each hormone. We investigated these possibilities by examining the expression of target genes of the three TFs shared between the BR, JA, SA and SL/KAR models: MYC2, MYC3 and WRKY33. The expression of many target genes of these TFs differed between hormone treatments (Fig. 2c; Extended Data Fig. 7b, c, d). For example, different clusters of MYC2 target genes were more highly expressed after JA treatment (cluster 10), SA and BR treatment (cluster 8) and SL/KAR treatment (cluster 1). This indicates that the functions or activity of shared TFs may differ between hormone regulatory pathways. Alternative possibilities are that unidentified competitor TFs regulate these same target genes, that different partner proteins may be recruited, or that the TFs themselves may be modified differently, under certain hormone conditions.

TFs can be differentially expressed in response to hormones, influencing TF abundance and activity (Chang et al., 2013; Zander et al., 2020). We determined that large and unique suites of TFs were differentially expressed in response to each hormone. In total, 849 (ABA), 542 (BR), 227 (ET), 304 (JA), 353 (SA) and 695 (SL/KAR) TFs were differentially expressed during the response to each hormone (Fig. 2d; Supplementary Table 7). A subset of these differentially expressed TFs was exclusive to one hormone, while only 29 TFs were shared between all hormones (Fig. 2d). In addition, a distinct pattern of enriched TF families was observed between the differentially expressed genes for each hormone (Fig. 2e; Supplementary Table 7). This, combined with the observations from the hormone transcriptional regulatory pathway models, indicates that dynamic remodeling of the transcriptome by ABA, BR, ET, JA, SA and SL/KAR involves large, hormone-specific suites of TFs even though hormones influence many overlapping growth and developmental processes. Each hormone recruits different combinations of TFs and activity of shared TFs may differ between hormones.

### Hub target genes are more highly responsive to hormones and are enriched in TFs

We investigated whether hub target genes exist within hormone transcriptional regulatory pathways and what their properties are. Hub targets are genes bound by many TFs (Heyndrickx et al., 2014). In plant transcriptional networks, unlike in animals, this high degree of binding is thought to be regulatory. Hub target genes may also exhibit different expression characteristics than non-hubs, being expressed under a wider range of conditions, likely as a result of being bound by many TFs (Heyndrickx et al., 2014). We identified the hub target genes in networks of 17 hormone TFs for which ChIP-seq data was available. We focused on this data type alone because it provides the most precise map of TF-target interactions (Fig. 3; Supplementary Table 8). A TF-target network was generated for each hormone because multiple hormone-specific ChIP-seq datasets were available for some TFs. Most genes (63.2% to 71.8% per hormone) were bound by more than one TF. The binding of target genes by multiple TFs was observed for all hormones, with some bound by as many as 12 TFs (Fig. 3a). We consequently defined hub target genes as genes bound by at least 7 TFs (Supplementary Table 8). By this threshold we identified 1,103 hub target genes across all hormones, compared with 15,203 non-hub target genes.

**Figure 3.**
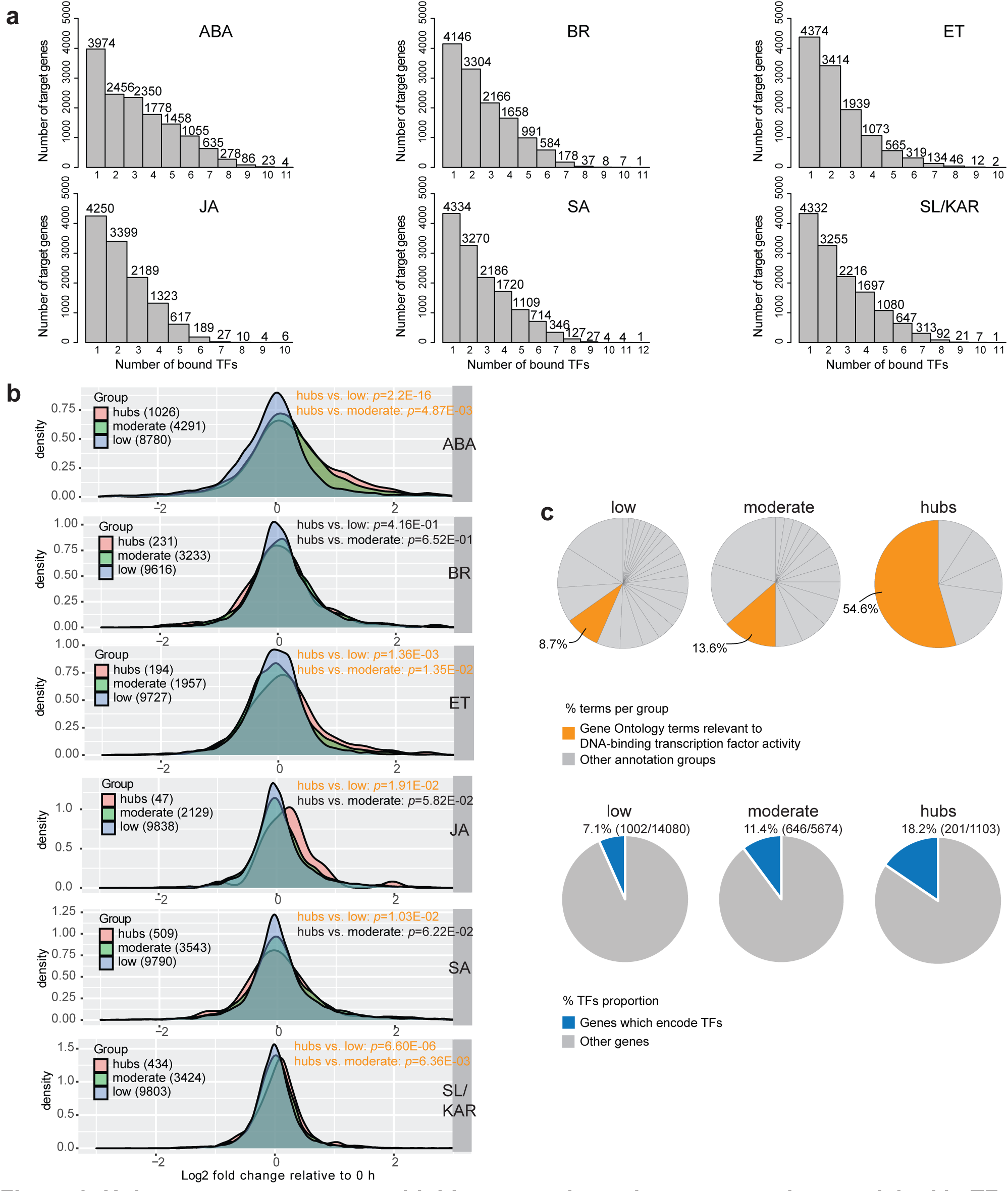
Hub target genes are more highly responsive to hormones and are enriched in TFs in hormone transcriptional networks. **a**, Plots show the number of TFs (x-axis) binding each target gene. Hub target genes were defined as genes bound by at least 7 TFs. **b**, Density plots show the differential expression for each target gene in three groups (low, genes bound by 1-3 TFs; moderate, genes bound by 4-6 TFs; hubs, genes bound by at least 7 TFs). The number of targets in each group are listed in parentheses. The x-axis is log2 fold change relative to 0 h. Each individual plot reports target gene expression for one of the six hormones. Significant difference in distribution is indicated by orange text (p-value < 0.05; two-sample Kolmogorov-Smirnov test). **c**, Pie charts in the upper panel show the percentage of DNA-binding transcription factor activity terms amongst all enriched gene ontology terms of the target genes. Individual pie charts present data for each group (low, moderate and hubs). The pie charts in the bottom panel give the proportion of genes encoding TFs in each group. The percentage and the numbers of TFs (in parentheses) in each group are listed at the top of corresponding pie chart.

The difference in the number of TFs that hub and non-hub target genes are bound by indicates that the expression of these two classes of genes may be regulated differently. The expression responses of hub and non-hub target genes to hormones differed, in accordance with this (Fig. 3b). Target genes were divided into three categories; low, moderate and hubs, bound by 1-3, 4-6 and 7 or more TFs, respectively. Differential expression for each target gene in the three categories was calculated, and plotted in density plots, then differences between distributions were assessed (Supplementary Table 8; Kolmogorov-Smirnov test, p-value < 0.05). Hub target genes were more highly differentially expressed in response to 5 of the 6 hormones than non-hub target genes (Fig. 3b). This indicates that the regulation of hub and non-hub target genes indeed differs. The increased differential expression may occur due to the additive action of the bound TFs bound at any gene. However, we highlight that TFs can have activator or repressive activity, leading to potentially conflicting influence upon the expression of bound genes.

Hub and non-hub target genes also differed in their annotated functional roles (Fig. 3c; Supplementary Table 8). Both hub and non-hub target genes were enriched in genes from hormone signaling pathways and genes with TF activity. However, enrichment for TF activity was greater amongst the hub target genes (Fig. 3c; low, 8.7%; moderate 13.6%; and hubs 54.6% of terms per group). Accordingly, more hub target genes encoded TFs than non-hub target genes.

Considered together, our findings indicate that a small proportion of genes in hormone transcriptional regulatory pathways are hub target genes, bound by many TFs. The existence of hub target genes may allow regulation to converge at certain genes, which presumably permits information from different signaling pathways to be integrated. Hub target genes are more strongly differentially expressed in response to hormones than non-hubs and are enriched for genes with TF activity. The hub target genes were similarly enriched for TFs in a network examining the expression of target genes of known flowering, circadian rhythm, and light response TFs (Heyndrickx et al., 2014). Given these similar features of the hormonal and flowering networks, it remains to be examined whether the TF activity of hub target genes is a more general principle of plant transcriptional networks.

### MAP kinases conduct cross-regulation between multiple hormones transcriptional regulatory pathways

Plant hormone signaling pathways do not operate in isolation from one another. Multiple contact points exist between different pathways, facilitating hormone cross-regulation (Altmann et al., 2020). We examined how cross-regulation between plant hormone signaling pathways influences TF activity and identified network components that may be responsible for cross-regulation. The most comprehensive current analysis of plant hormone signaling cross-regulation is a large-scale protein-protein interaction network (Altmann et al., 2020). We extended upon this by connecting hormone signaling protein-protein interactions to TF-gene interactions and gene expression. To do so, we generated an integrated transcriptional cross-regulation model by overlaying the individual hormone transcriptional regulatory models (Fig. 1a; 4a; Supplementary Table 5). The integrated model was composed of 291 individual genes, 23 of which were shared by at least 4 hormones (Fig. 4a, b; Supplementary Table 5). These 23 shared genes were 13 TFs from different families, 8 MITOGEN-ACTIVATED PROTEIN KINASES (MPKs) and the genes TPL (AT1G15750) and POLYUBIQUITIN 3 (UBQ3, AT5G03240). The 23 genes were significantly enriched for signal transduction functions (Fig. 4c; p-value < 0.01). We infer that these genes are likely nodes of cross-regulation in a broad, multi-hormone regulatory network.

**Figure 4.**
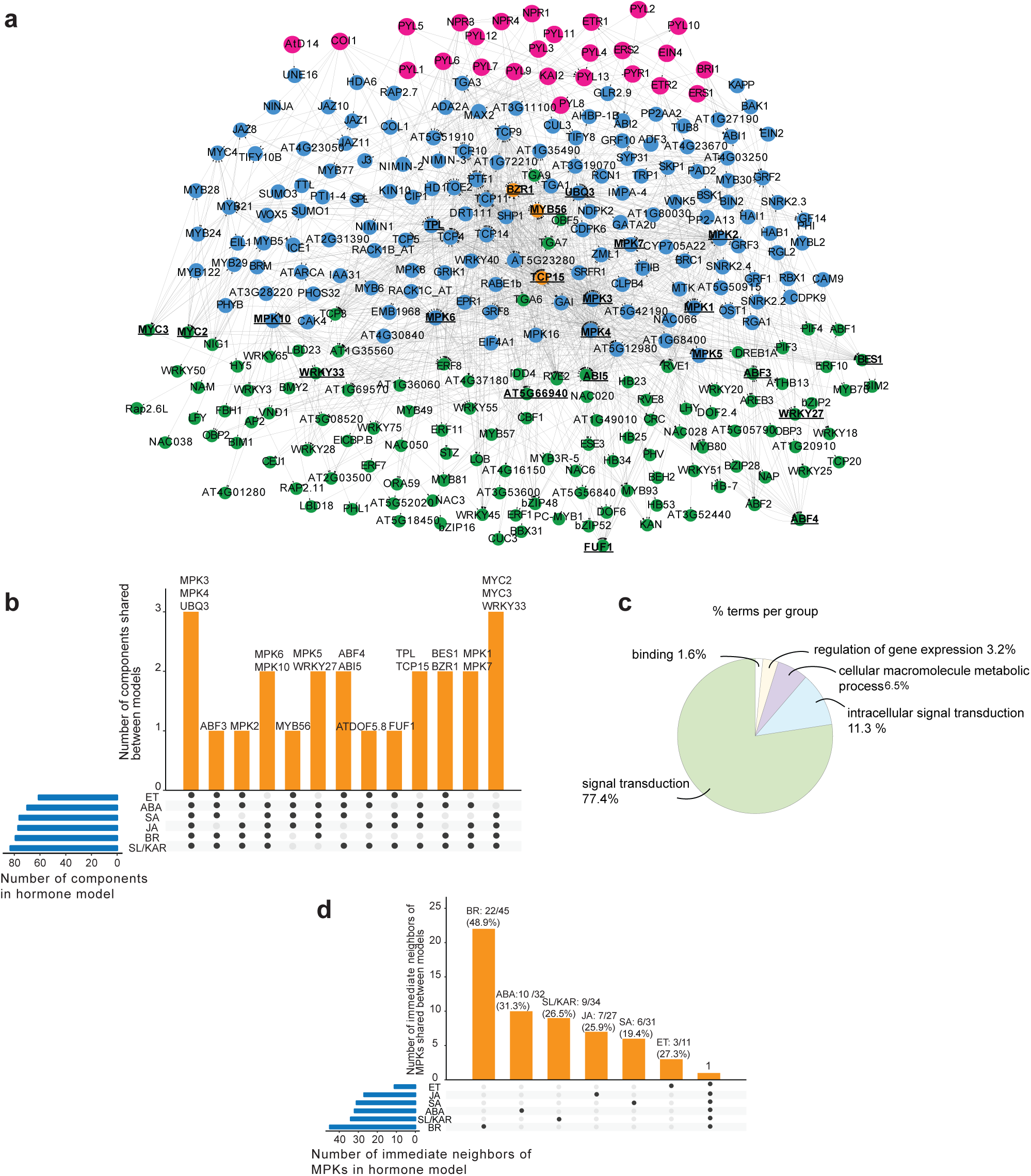
MPKs are convergence nodes in the integrated multi-hormone cross-regulation model. **a**, The multi-hormone cross-regulation network was built by integrating models of each hormone. Magenta nodes: upstream proteins given as hormone receptors. Blue nodes: predicted signaling proteins. Green nodes: active TFs responsible for transcriptional changes. Orange nodes: proteins that have both signaling and active TFs roles. The proteins shared by at least 4 hormone pathways are in black bold text and have underlined names. **b**, Twenty-three proteins are shared by at least 4 hormone signaling pathways and are putative hormone cross-regulation nodes. These proteins were enriched in MPKs (8/23). **c**, Pie chart shows the functional groups of enriched (p-value < 0.05) gene ontology terms of 23 proteins. The listed group name is the term has highest significance in its functional group. The percentages of the enriched terms in each group amongst all enriched gene ontology terms of 23 proteins are listed following the group names. **d**, Unique and shared immediate neighbors of MPKs in each individual hormone model. The numbers and percentages at the top of each column indicate what proportion the unique immediate neighbors are of the total immediate neighbors.

The 23 predicted multi-hormone cross-regulation genes included TFs directly and indirectly associated with hormone signaling. Seven of thirteen TFs (53.8%) were known regulators of hormone responses (Supplementary Table 5). These were ABSCISIC ACID RESPONSIVE ELEMENTS-BINDING FACTOR 3 (ABF3), ABF4, ABA INSENSITIVE 5 (ABI5), MYC2, MYC3, BRI1-EMS-SUPPRESSOR 1 (BES1) and BRASSINAZOLE-RESISTANT 1 (BZR1), associated with the ABA, JA and BR signaling pathways (Choi et al., 2000; He et al., 2005; Li and Deng, 2005; Fujita et al., 2013; Kazan and Manners, 2013; Salazar-Henao et al., 2016; Skubacz et al., 2016; Hickman et al., 2017; Ibanez et al., 2018; Ju et al., 2019; Chen et al., 2020; Zander et al., 2020). In addition, two of these TFs have defined roles in cross-regulation between a small number of hormones (ABI5 - ABA and JA, ET; MYC2 - JA and ET, ABA, SA) (Abe et al., 2003; Wild et al., 2012; Zhang et al., 2014; Ju et al., 2019). The remaining TFs were not characterized as directly involved in hormone signaling but had roles in processes associated with hormones, such as plant defense, abiotic stress responses and flowering (Zheng et al., 2006; Mukhtar et al., 2008; Pandey and Somssich, 2009; Chen et al., 2015; He et al., 2015). These shared TFs provide a potential mechanism for cross-regulation of gene expression between multiple hormone signaling pathways.

The large number of MPKs present amongst the 23 predicted multi-hormone cross-regulation genes was notable (8/23 genes, Fig. 4b; Supplementary Table 5). Protein phosphorylation cascades transduce developmental and environmental signals and regulate cell functions *via* MPKs (Cristina et al., 2010; Bigeard and Hirt, 2018; Jagodzik et al., 2018). MPKs often occupy core positions in signal transduction pathways, receiving information from several upstream inputs. They then phosphorylate downstream proteins, frequently including TFs, thereby regulating their activity. These properties would make them extremely suitable as central components for multi-hormone cross-regulation. TFs are a large proportion of the immediate (first-degree) neighbors of MPKs in our model (73.5%, 75/102; Supplementary Table 5). This is consistent with the previous results of a MPK target network (Popescu et al., 2009). Many of these immediate neighbors were unique to individual hormones (ABA, 10/32, 31.3%; BR, 22/45, 48.9%; ET, 3/11, 27.3%; JA, 7/27, 25.9%; SA, 6/31, 19.4%; SL/KAR, 9/34,

26.5%; Fig. 4d; Supplementary Table 5). This indicates that, despite being shared between hormone pathways, the MPKs likely target different downstream proteins dependent upon the hormone they respond to.

### Mutation of *MPK6* broadly affects the hormone-responsive proteome, phosphoproteome and transcriptome

We next validated a candidate multi-hormone cross-regulation node, *MPK6*, as having effects consistent with a central role in the network. We did so by examining the broad-scale influence of *mpk6* mutation on hormone-responsive transcription, protein abundance and protein phosphorylation. We generated transcriptomic, proteomic and phosphoproteomic data using wild-type (WT; Columbia-0: Col-0) and *mpk6* mutant etiolated seedlings following hormone or mock treatment (ABA; ET, using the ET precursor 1-aminocyclopropane-1-carboxylic acid, ACC, which stimulates ET signaling; JA, SA) for 1 h, then compared the responses of WT and *mpk6* mutant seedlings (Fig. 5; Extended Data Fig. 8a, b, c, d; Supplementary Table 9, 10, 11, 12).

**Figure 5.**
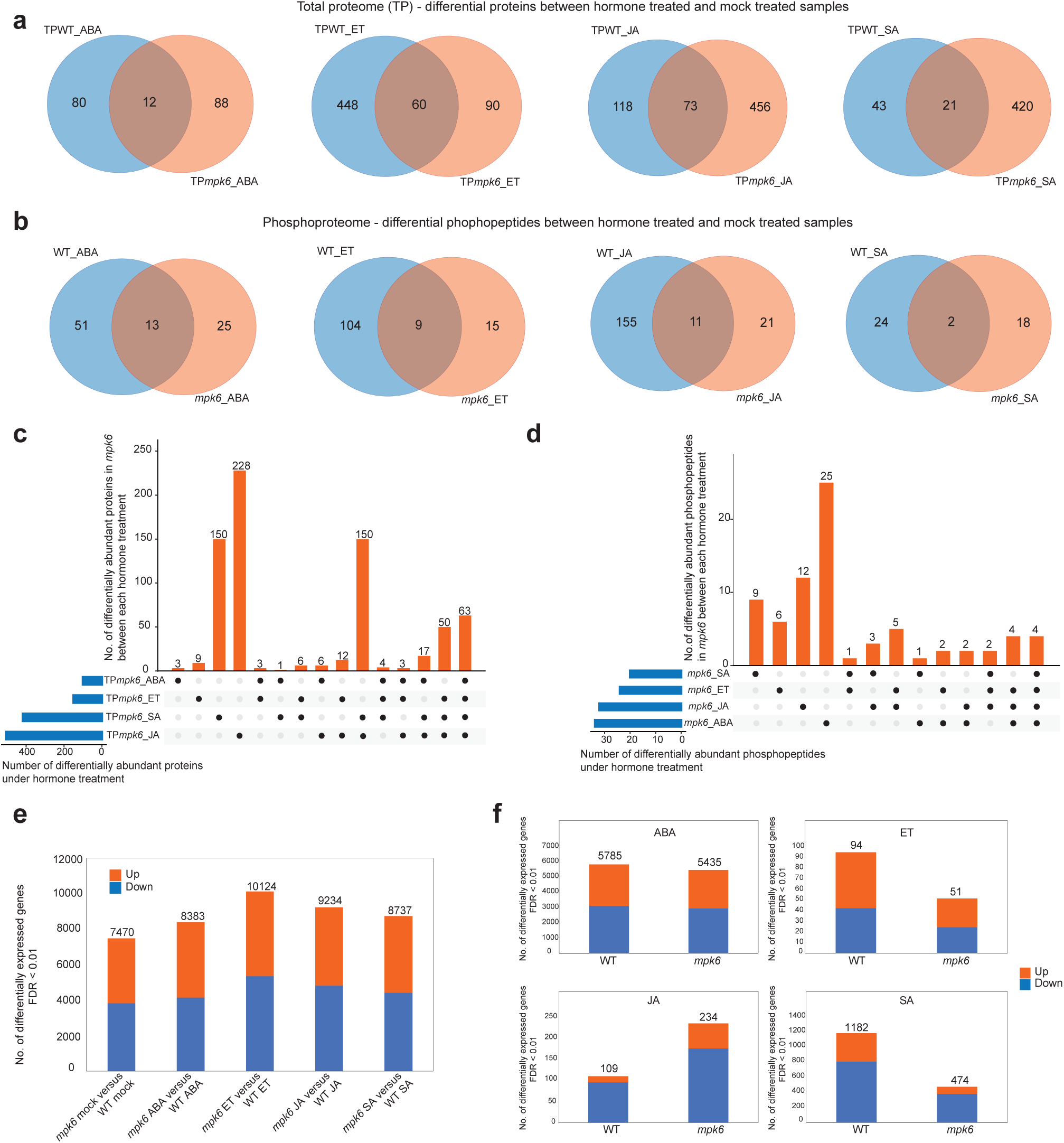
Mutation of *MPK6* broadly affects the hormone-responsive proteome, phosphoproteome and transcriptome. **a**, Unique and shared differentially abundant proteins between WT and *mpk6* seedlings following treatment with hormones (ABA, ET, JA, SA). For each hormone, proteomes of WT or *mpk6* seedlings after hormone treatment were compared to their respective mock treated samples and differentially abundant proteins identified (p-value < 0.05 & fold change > 1.1) from total proteome analysis. Venn diagrams represent the overlap of these differentially abundant proteins between genotypes. **b**, Unique and shared differentially abundant phosphopepetides between WT and *mpk6* seedlings following treatment with hormones. Comparisons were conducted as for proteomes but using phosphoproteomic data. **c**, Unique and shared differentially abundant proteins in *mpk6* between each hormone treatment. **d**, Unique and shared differentially abundant phosphopeptides in *mpk6* between each hormone treatment. **e**, Numbers of significantly differentially expressed genes (edgeR; FDR < 0.01) between *mpk6* and WT seedlings after mock treatment and each hormone treatment. The numbers of significantly up-regulated and down-regulated genes are indicated separately by the orange (up) and blue (down) sections of bars. **f**, Total number of significantly differentially expressed genes (edgeR; FDR < 0.01) detected in comparisons between hormone treated WT and *mpk6* seedlings and mock treated samples.

The *MPK6* mutation had broad effects on protein abundance and phosphorylation following hormone treatment, indicating that MPK6 has a role in responses to all four hormones. A considerable number of proteins and phosphopeptides were significantly differentially abundant in hormone-treated WT and *mpk6* etiolated seedlings compared with their respective mock treated samples (64 – 529 proteins, 20 – 166 phosphopeptides, p-value < 0.05 & fold change > 1.1; Extended Data Fig. 8e; Supplementary Table 9, 10). The majority of these hormone-responsive proteins and phosphopeptides were unique to either WT or *mpk6* (WT proteins: 80/92, 87.0%, ABA; 448/508, 88.2%, ET; 118/191, 61.8%, JA; 43/64, 67.2%, SA; *mpk6* proteins: 88/100, 88%, ABA; 90/150, 60%, ET; 456/529, 86.2%, JA; 420/441, 95.2%, SA; WT phosphopeptides: 51/64, 79.7%, ABA; 104/113, 92.0%, ET; 155/166, 93.4%, JA; 24/26, 92.3%, SA; *mpk6* phosphopeptides: 25/38, 65.8%, ABA; 15/24, 62.5%, ET; 21/33, 65.6%, JA; 18/20, 90.0%, SA; Fig. 5a, b; Supplementary Table 11). Furthermore, *mpk6* altered the abundance of different proteins and phosphoproteins between each hormone response (Fig. 5c, d). These results are consistent with MPK6 having a role in multiple hormone signaling pathways and that the role of MPK6 differs between hormones.

Mutation of *MPK6* changed the transcriptional response of seedlings to hormone treatment. The transcriptomes of *mpk6* mutant seedlings differed substantially from transcriptomes of WT seedlings, both in the absence of hormone and after hormone treatment (numbers of differentially expressed genes; 7,470, *mpk6* mock *v*. WT mock; 8,383, *mpk6* ABA *v*. WT ABA; 10,124, *mpk6* ET *v*. WT ET; 9,234, *mpk6* JA *v*. WT JA; 8,737, *mpk6* SA *v*. WT SA; Fig. 5e; Supplementary Table 12). However, *mpk6* plants did still respond to hormone treatment, with comparable numbers of transcripts being differentially expressed following hormone treatment in *mpk6* and WT relative to their respective mock treated samples (Fig. 5f; Supplementary Table 12). Many TFs were differentially expressed between *mpk6* and WT seedlings after hormone treatment (Extended Data Fig. 8f). These results indicate that the *mpk6* mutation disrupts, but does not eliminate, hormone-responsive changes to the transcriptome.

These findings indicate that loss of functional MPK6 extensively remodels the total proteome, phosphoproteome and transcriptome of multiple hormone responses, and that it influences each hormone response differently. This is consistent with a proposed role of MPK6 as a central regulator of multi-hormone cross-regulation, potentially affecting the activity of downstream TFs in hormone gene regulatory networks.

### Alternative splicing is a core component of hormone responses

Alternative splicing contributes to the reprogramming of gene expression, changing the functional composition of proteins expressed from individual genes (Narsai et al., 2017; Calixto et al., 2018). JA responses include alternative splicing, but the influence of alternative splicing on broader hormone responses is not characterized (Chung et al., 2010; Moreno et al., 2013; Zander et al., 2020). To examine this, we identified genes whose transcripts were differentially alternatively spliced following each hormone treatment. We analyzed the time-series RNA-seq data at transcript-level for all hormones except ET; the ET sequence read length was too short for this analysis. There were 1,155 (ABA), 1,016 (BR), 415 (JA), 613 (SA), and 797 (SL/KAR) genes whose transcripts were differentially alternatively spliced during the response to each hormone (Fig. 6a; Supplementary Table 13). Amongst these, 818 (ABA), 376 (BR), 96 (JA), 153 (SA) and 411 (SL/KAR) were also differentially expressed at gene-level, which indicates that many genes are regulated by both transcription and alternative splicing (Fig. 6b). However, substantial numbers of genes were not differentially expressed, and consequently regulated only by alternative splicing (337, 29.2%, ABA; 640, 63.0%, BR; 319, 76.9%, JA; 460, 75.0%, SA; 386, 48.4%, SL/KAR; Fig. 6b). The three most abundant types of events amongst hormone responsive differentially alternative spliced transcripts were intron retention (36.7 - 38.7%), alternative 3’ splice sites (28.0-29.1%) and alternative 5’ splice sites (21.4 - 21.7%) (Extended Data Fig. 9a; Supplementary Table 13). This was common across all hormones and is comparable with alternative splicing during cold responses (Calixto et al., 2018). Furthermore, a large proportion of differentially alternatively spliced genes was unique to individual hormones (Extended Data Fig. 9b). These results demonstrate that alternative splicing is a general component of plant hormone signaling.

**Figure 6.**
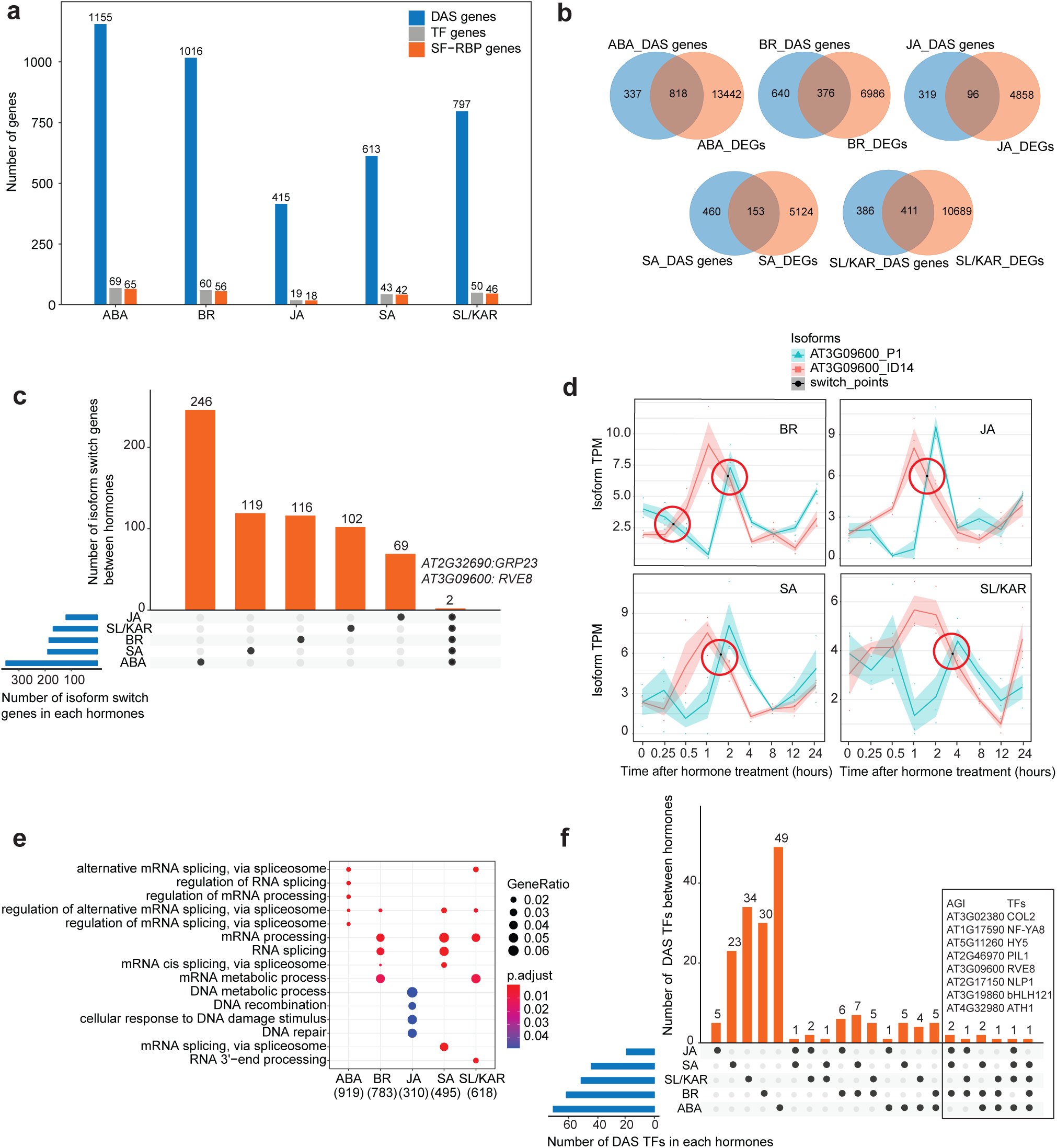
Alternative splicing is a core component of hormone responses. **a**, The numbers of significant differentially alternative spliced (DAS) genes (FDR < 0.05), and the number of genes encoding TFs, and splicing factors and RNA binding proteins (SF-RBPs) amongst the DAS genes following each hormone treatment. **b**, Overlap between DAS genes and differentially expressed genes (DEGs) for each hormone. **c**, Number of genes that exhibit isoform switching between hormones. Isoform switch event describes the splicing phenomenon whereby the relative abundance of two transcript isoforms from a single gene reverse following hormone treatment. The plot shows how many are unique to a hormone and how many are shared between all five hormones analyzed. Two genes are shared by 5 hormones, whose gene ID and names are labelled at the top of respective columns. **d**, Example isoform switch events for RVE8 (two different isoforms, P1 vs. ID14) for four hormone responses. The isoform switch points detected by TSIS are indicated with red circles. **e**, Top five significantly enriched (p.adjust < 0.05) gene ontology terms amongst the DAS genes upon each hormone treatment. **f**, DAS TFs that are unique and shared between the five hormone responses analyzed.

Isoform switching is a phenomenon whereby the relative abundance of two transcript isoforms from a single gene reverse following a stimulus (Guo et al., 2017). Such events change the dominant form of the transcript present and the structure of the subsequent mature protein, which can influence cellular processes (Chung et al., 2009). Isoform switching occurs during the JA response in etiolated Arabidopsis seedlings, but its contribution to other hormone responses is unknown (Zander et al., 2020). We found 350 (ABA), 185 (BR), 120 (JA), 190 (SA) and 169 (SL/KAR) genes underwent isoform switching (Supplementary Table 14). Almost all of these isoform switching events involved at least one protein-coding transcript isoform (456, 96.6%, ABA; 229, 97.0%, BR; 132, 96.4%, JA; 241, 98.0%, SA; 190, 95.0%, SL/KAR; Supplementary Table 14). This indicates that the events can potentially change the function of the proteins expressed in a hormone response. The majority of isoform switching genes differed between hormones (Fig. 6c). Transcripts of two genes underwent isoform switching in response to all 5 hormones (*AT3G09600*, also known as *REVEILLE 8* and *RVE8*; *AT2G32690*, also known as *GLYCINE-RICH PROTEIN 23* and *GRP23*). However, different pairs of *GRP23* isoforms were affected across the five hormones. *RVE8* isoform switching was dominated by one pair across four hormones, but the switch time point differed (Fig. 6d). These results indicate that isoform switching is a common feature of hormone responses.

The functional influence of alternative splicing on plant hormone responses is also unknown. Two molecular characteristics were notable amongst the hormone-responsive differentially alternatively spliced genes. First, mRNA splicing-related functions were enriched for 4 of 5 hormones (Fig. 6e; Supplementary Table 13). Accordingly, 4.3 - 6.9% of differentially alternatively spliced genes were splicing factors, RNA binding proteins (SF-RBPs) or spliceosome proteins. Second, transcripts encoding many TFs were differentially alternative spliced during hormone responses (Fig. 6a; Supplementary Table 13). The majority of these were alternatively spliced uniquely in response to a single hormone, except for JA-responsive TFs (unique splicing events; 71.0%, ABA; 50.0%, BR; 26.3%, JA; 53.5%, SA; 68.0%, SL/KAR; Fig. 6f; Supplementary Table 13), indicating that individual hormones regulate distinct sets of TFs through alternative splicing. Nevertheless, a small number of common alternatively spliced TFs did exist, with eight TFs shared by at least three hormones. Amongst these, published data connected to hormone signaling existed only for ELONGATED HYPOCOTYL5 (HY5) (Hamasaki et al., 2020; Ortigosa et al., 2020). These findings indicate that hormone responses use alternative splicing to further diversify the transcriptome by influencing TFs and the alternative splicing machinery itself.

### Alternative splicing contributes to early hormone signaling responses independent of differential gene expression

Alternative splicing is co-transcriptional but can occur without differential gene expression (Marquez et al., 2012). We consequently examined the relative dynamics of alternative splicing and differential expression during hormone responses, to better understand their relative contributions to transcriptome reprogramming. We determined the time at which each gene or transcript was first significantly differentially expressed or alternatively spliced relative to 0 h (Fig. 7a). The temporal dynamics of differential expression and alternative splicing differed, with alternative splicing appearing to be more frequent at early time points than differential expression. To examine this more closely we plotted the relative proportion of alternatively spliced and differentially expressed genes across all time points (Fig. 7b). The proportional contribution of alternative splicing to transcriptome remodeling was greatest at early time points for all hormones except BR. These responses occurred within the first 15 mins to 1 h after hormone treatment. Consequently, it is likely that alternative splicing has an important role in rapid responses to hormone signaling, acting independently of differential gene expression.

**Figure 7.**
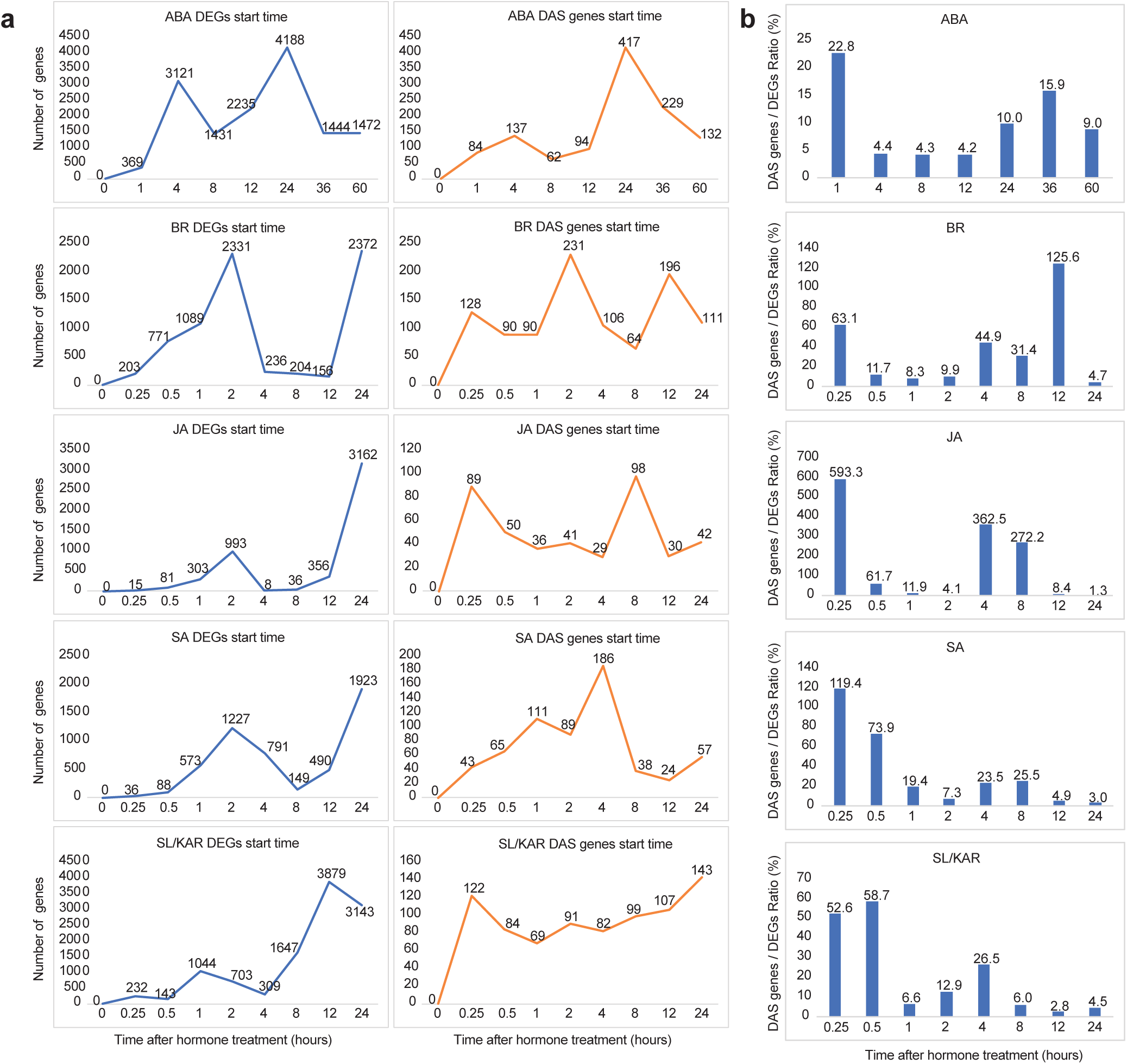
The relative dynamics of differential expression and alternative splicing in hormone responses. **a**, The number of genes or transcripts first significantly differentially expressed or alternatively spliced relative to 0 h at each time point. **b**, Plots show the relative proportion of differentially alternative spliced (DAS) genes and differentially expressed genes (DEGs) across all time points in each hormone dataset.

## Discussion

Plant hormones do not operate in isolation. Rather, they form a large network to optimize plant growth and development (Altmann et al., 2020). In this study, we aimed to determine the extent of cross-regulation in a network of 6 hormones and to understand how this was related to gene expression. We reconstructed a dynamic signaling gene regulatory network model of hormone cross-regulation in etiolated Arabidopsis seedlings. This was achieved by integrating time-series transcriptomics following ABA, ET, JA, SA, BR or SL/KAR treatment with genome-wide target maps for hundreds of TFs and large-scale protein-protein interaction maps. Our major finding from these analyses is that hormone cross-regulation occurs at multiple levels, spanning signal transduction, TF activity and gene expression. This integrated view extends our knowledge of the scale and mechanisms of hormone cross-regulation. It also increases our understanding of the fundamental principles of plant transcriptional regulation during hormone responses. The framework we have developed here to characterize regulatory networks in environmental responses and development can be applied broadly to plant biology and beyond.

Our study illustrates that different hormones employ different combinations of TFs for transcriptional regulation, but a small number of TFs are shared between hormones. The functions of shared TFs may differ between hormone regulatory pathways as their target genes may vary, or if they have a differential function as an activator or repressor. We identified that hub target genes bound by multiple TFs exist in hormone responses, allowing regulation to converge at certain genes. Furthermore, hub target genes exhibit stronger differential expression in response to hormones and greater enrichment for TF activity than non-hubs. These properties are consistent with the properties reported of hub target genes in a network of Arabidopsis flowering and light regulation, as well as a transcriptional regulatory network of maize leaf (Heyndrickx et al., 2014; Tu et al., 2020). Considering these together, our findings demonstrate that the regulatory activity of TFs during hormone responses is complex. The results also provide evidence that the TF activity of hub target genes may be a general principle of plant regulatory networks.

In this study we predicted that a group of MPKs may have a central role in hormone cross-regulation. Their biochemical properties mean they are very well-suited to this. MPK cascades are a highly conserved feature of the signaling pathways that integrate environmental signals into rapid cellular responses (Cristina et al., 2010; Raja et al., 2017; Bigeard and Hirt, 2018; Jagodzik et al., 2018). The roles of some of these MPKs in single or interactions between pairs of hormone signaling pathways have been well studied (Jagodzik et al., 2018). Our study differs importantly, however, because we demonstrate that MPKs may act as convergence points for cross-regulation within a network of multiple hormones. In addition, we observed that MPKs have many immediate neighbors that differ between specific hormone responses, which may indicate that they drive different responses between hormones. However, the mechanisms of how MPKs select and phosphorylate different downstream substrates between hormone signaling pathways are unknown. This may be programmed through modifiers such as the SMALL UBIQUITIN-LIKE MODIFIER (SUMO) interaction motif of MPK3/6, which enables MPK3/6 to differentially select and phosphorylate substrates (Verma et al., 2021). Moving forward it will be important to functionally validate the predicted roles of MPKs in hormone responses by comprehensively identifying the interactors and targets of these MPKs. This will allow us to better understand how plants process multiple hormone signals through MPKs to adapt to diverse environmental conditions.

Alternative splicing occurred within the early time points of hormone responses and made its largest contribution within this time window. This suggests alternative splicing acts independent of differential gene expression to some extent, which differs from plant responses to cold where alternative splicing accompanies the major transcriptional changes (Calixto et al., 2018). Hundreds of TFs, splicing factors and RNA binding proteins were alternatively spliced, all of which would act to further diversify the transcriptome and, presumably, the proteome. These features were common across all hormones. This may indicate that RNA splicing is an important component of hormone signaling mechanisms. The digestion strategy we used in our proteomic analyses did not permit detection of proteins arising from alternative splicing events due to the sequence preferences of trypsin, which targets lysine and arginine-coding triplets that tend to be evolutionarily conserved at intron-exon boundaries, but in future this might be examined using alternative approaches (Wang et al., 2018).

Overall, our study provides a broad view of how multiple hormone signals interact as a network to cross-regulate gene expression. This provides a framework for interrogating temporal dynamics of the hormone-responsive transcriptome. Future studies might consider the relationship between spatial and temporal regulation of gene expression in dynamic responses.

## Methods

### Plant materials, growth conditions and hormone treatments

Three day old etiolated Arabidopsis seedlings of Col-0 background were used for all RNA-seq, ChIP-seq, and (phospho)proteomic experiments. The transgenic lines Col-0 ANAC055::ANAC055-YPet, Col-0 BES1::BES1-YPet, Col-0 EDF1::EDF1-YPet, Col-0 EDF2::EDF2-YPet, Col-0 EDF3::EDF3-YPet, Col-0 EIN3::EIN3-YPet, Col-0 ERF1::ERF1-YPet, Col-0 MYC2::MYC2-YPet, Col-0 MYC3::MYC3-YPet, Col-0 OBP2::OBP2-YPet, Col-0 RAP2.6L::RAP2.6L-YPet, Col-0 STZ::STZ-YPet, Col-0 TCP3::TCP3-YPet and Col-0 TGA5::TGA5-YPet, were generated by recombineering, as previously described (Zander et al., 2020).

Seeds were sterilized with bleach and sown on Murashige and Skoog (cat#LSP03, Caisson) media pH 5.7, containing 1% sucrose and 1.8% agar. After stratification (three days dark at 4°C), seeds were exposed to light at room temperature for 2 hours to induce germination, then grown in the dark at 22°C for three days. Etiolated seedlings were subsequently treated with hormones.

For time series RNA-seq experiments, BR, SA and SL/KAR treatments were applied by spraying the plants until run-off occurred then samples were harvested at each time point. BR treatment was conducted using 10 nM epibrassinolide (Sigma, E1641), SA treatment using 0.5 mM SA (Sigma, S7401), and SL/KAR treatment using 5 μM rac-GR24 (Chiralix, Nijmegen, The Netherlands). For ChIP-seq experiments, BR and SA treatments were as described above. JA and ET treatments were performed as previously described (Chang et al., 2013; Zander et al., 2020).

*mpk6* mutant seeds (*mpk6_4*, SALK_062471C) were obtained from the Arabidopsis Biological Resource Centre (ABRC). This Col-0 background mpk6 allele has been extensively characterized and validated as a total knockout of *mpk6* expression (Bush and Krysan, 2007; Merkouropoulos et al., 2008; Li et al., 2017). Prior to use, the genotype of this line was confirmed by conducting PCR for the SALK T-DNA insert. Seeds were stratified in water for 3 days at 4°C in the dark then sown on soil (standard potting mix, Van Schaik’s BioGro, Australia). Plants were grown to maturity in a controlled environment room with a 16/8-hour light/dark cycle, 22°C/19°C (light/dark), 55% relative humidity and 120 μmol m−2 s−1 light intensity. Leaf samples were harvested by snap-freezing in liquid N2 and stored at -80°C. Then frozen samples were ground using a TissueLyser II (QIAGEN) and extracted with a fast genomic DNA extraction protocol (Kasajima et al., 2004). PCR was performed using PCR master mix (Thermo Fisher Scientific, M0486L) according to manufacturer’s instructions. Primer pairs for genotyping were designed using online website tool http://signal.salk.edu/tdnaprimers.2.html. Left genomic primer (LP): GTCCAGGGAAGAGTGGCTTAC; Right genomic primer (RP): GCAGTTCGGCTATGAATTCTG. Two paired reactions were set up for the following PCR reactions; LP plus RP for detecting the presence of a WT copy of the gene, and left border primer of the T-DNA insertion (LBb1.3: ATTTTGCCGATTTCGGAAC) plus RP for detecting the T-DNA/genomic DNA junction sequence. Homozygosity of the line was confirmed if it exhibited the pattern of no product for WT copy but positive for the T-DNA product. PCR was conducted on the T100 Thermal Cycler PCR system (BIO-RAD) with PCR conditions as follows: 94 °C for 30 sec, and 30 cycles at 94 °C for 30 sec / 55 °C for 30 sec, 68 °C for 1 min 10 sec, then 68°C for 5 min. PCR products were analaysed by gel electrophoresis and imaged on the Gel DocTM XR+ system (BIO-RAD). Only seeds from homozygous plants were used in subsequent experiments.

For *mpk6* RNA-seq, proteomic and phosphoproteomic analyses, WT and *mpk6* seeds were sterilized using chlorine gas in a desiccator for 3 hours. The seeds were then plated on ½ MS media supplemented with 1% sucrose and 1.5% agar, pH 5.7. The plates were wrapped, and seeds were stratified for 3 days at 4 °C, then brought to 22°C under the light for 2 hours before wrapping up again with aluminium foil and growing for another 3 days. Plates were then taken to a dark room that was illuminated with green light (Lee Sheet 736 Twickenham Green). Three day old etiolated seedlings were sprayed with hormones (ACC, 10 µM; SA, 0.5 mM; 10 nM, ABA 10 µM) or treated with JA as described above. Hormone treatments lasted for an hour, then seedlings were harvested and snap-frozen in liquid nitrogen.

### ChIP-seq data generation

All ChIP-seq sequencing data was generated using biologically independent replicate experiments: ANAC055 (air, n=4; ET, n=4), ANAC055 (air, n=3; JA, n=4), BES1 (air, n=4; ET, n=4), BES1 (BR, n=2), EDF1 (air, n=3; ET, n = 3), EDF2 (air, n=3; ET, n=4), EDF2 (JA, n=2), EDF3 (air, n = 3; ET, n = 3), EIN3 (air, n = 3; ET, n= 4), EIN3 (JA, n=2), ERF1 (JA, n=3), MYC2 (air, n=3; ET, n = 3), MYC2 (air, n=3; JA, n=3), MYC3 (air, n=3; ET, n=3), MYC3 (air, n=4; JA, n=4), OBP2 (air, n=2; ET, n=2), OBP2 (air, n=2; JA, n=3), RAP2.6L (air, n=3; ET, n=3), RAP2.6L (air, n=2; JA, n=2), STZ (air, n=3; JA, n=4), STZ (air, n=2; ET, n=1), TCP3 (air, n=4; ET, n=4), TCP3 (air, n=3; JA, n=3), TCP3 (BR, n=1), TGA5 (air, n=2; ET, n=2), TGA5 (SA, n=1). The ChIP-seq data for JA-treated ANAC055, MYC2, MYC3 and STZ, and the ChIP-seq data for air-treated STZ has been reported in our previous publication (Zander et al., 2020). The remaining ChIP-seq data was generated by this study.

Seedlings were treated with BR, ET, JA, SA or air for 2 hours then collected and snap-frozen in liquid nitrogen. Chromatin preparation and immunoprecipitation were performed as previously described (Zander et al., 2020). A goat anti-GFP antibody (supplied by D. Dreschel, Max Planck Institute of Molecular Cell Biology and Genetics) was used and mock immunoprecipitations were conducted using whole goat IgG (005–000–003, Jackson ImmunoResearch). Immunoprecipitated DNA was used to prepare sequencing libraries. Libraries were sequenced on an Illumina HiSeq 2500 per manufacturer’s instructions (Illumina).

The raw ChIP-seq data for three ABFs (ABF1, ABF3, ABF4) under air and ABA treatment was downloaded from (Song et al., 2016). They were re-analyzed using the uniform workflow described in the following ChIP-seq analyses section.

### ChIP-seq data analyses

We developed an analysis workflow to process all raw fastq data in a uniform and standardized manner to enable integration and comparison. Fastq files were trimmed using Trimglore V0.4.4 then trimmed reads were mapped to the Arabidopsis TAIR10 genome using Bowtie2 V2.2.9 (Langmead and Salzberg, 2012). The mapped reads were filtered with MAPQ > 10 using samtools V1.3.1 to restrict the number of reads mapping to multiple locations in the genome (Li et al., 2009). Filtered reads were used for all the subsequent analyses. PhantomPeakQualTools v.2.0 was used to assess ChIP-seq experiment quality after read mapping by determining the normalized strand cross correlation (NSC) and relative strand cross correlation (RSC) of each alignment bam file.

MACS V2.1.0 was used to identify the peaks for all replicates by comparison with mock IP of wild-type Col-0 (default parameters except –g 1.19e8 and –q 0.05) (Zhang et al., 2008). Mapped reads and peak locations were visualized using JBrowse (Buels et al., 2016). Only peaks with a q-value >= 1^e-15^ were used in following analyses. Furthermore, only replicates with more than 50 peaks were retained. The total numbers of biological replicates retained for peak annotation were: ANAC055 (air, n=3; ET, n=3), ANAC055 (air, n=2; JA, n=3), BES1 (air, n=2; ET, n=1), BES1 (BR, n=1), EDF1 (air, n=1; ET, n =2), EDF2 (air, n=2; ET, n=3), EDF2 (JA, n=2), EDF3 (air, n =1; ET, n =3), EIN3 (air, n =2; ET, n=3), EIN3 (JA, n=2), ERF1 (JA, n=3), MYC2 (air, n=3; ET, n =2), MYC2 (air, n=3; JA, n=3), MYC3 (air, n=3; ET, n=3), MYC3 (air, n=3; JA, n=3), OBP2 (air, n=2; ET, n=2), OBP2 (air, n=1; JA, n=2), RAP2.6L (air, n=3; ET, n=2), RAP2.6L (air, n=1; JA, n=1), STZ (air, n=1; JA, n=2), STZ (ET, n=1), TCP3 (air, n=3; ET, n=3), TCP3 (air, n=2; JA, n=2), TCP3 (BR, n=1), TGA5 (air, n=2; ET, n=2), TGA5 (SA, n=1). In general, there were at least two biological replicates for each ChIP-seq sample (35/45, 77.8%).

For each TF, peaks that had at least 50% intersection in at least two independent biological replicates were merged using bedtools V2.26.0 and retained, with all other peaks eliminated (Quinlan and Hall, 2010). Peaks were associated to their nearest genes as annotated in the TAIR10 using R package ChIPpeakAnno with default parameters (Zhu et al., 2010).

### RNA isolation and library preparation

For time-series RNA-seq experiments, total RNA was isolated from liquid nitrogen ground whole etiolated seedlings using the RNeasy Plant Kit (Qiagen, CA, USA). cDNA libraries were constructed using the Illumina TruSeq Total RNA Sample Prep Kit (Illumina, CA, USA) as per manufacturer’s instructions. Single-end reads were generated by the HiSeq 2500 Sequencing System (Illumina). For *mpk6* validation RNA-seq experiments, RNA extractions were carried out using Sigma Spectrum Plant Total RNA Kit, supplemented with Sigma On-Column DNase I Digestion Set, according to the manufacturer’s instructions. Two µg of RNA was used for RNA sequencing library construction, using Illumina TruSeq Stranded Total RNA kit. RNA-seq libraries were then pooled into one and sequenced using a NovaSeq S1 Flow-cell, 100 bp single-end reaction.

### RNA-seq analyses

FastQC V0.11.5 was used to perform quality control. Trimglore V0.4.4 (https://www.bioinformatics.babraham.ac.uk/projects/) was used to remove low-quality reads and adapters from raw RNA-seq reads. Trimmed reads of the ET time series transcriptome data were mapped onto the Arabidopsis genome with the Araport11 annotation using HiSat2 V2.0.5 (Kim et al., 2015). Read counting in genome features was performed using Htseq V0.8.0 (Anders et al., 2015). This different process was necessary because the ET RNA-seq data were from color-space sequencing (ABI SoLID platform). For the other five hormone time series transcriptome datasets (ABA, BR, JA, SA, SL/KAR) and the RNA-seq datasets generated for validation (ABA, ET, JA, SA), quantification of transcripts was performed using Salmon v0.8.1 in conjunction with AtRTD2-QUASI reference transcriptome (Zhang et al., 2017). A quasi-mapping-based index was built using an auxiliary k-mer hash over k-mers of length 31 (k=31). Salmon parameters were kept default for quantification except that fragment-level GC biases (“–gcBias”) correction was turned on. The Tximport pipeline was used to summarize transcript-level abundance to gene-level abundance.

Differentially expressed genes in time-series and validation experiment RNA-seq were identified using edgeR 3.28.1 with quasi-likelihood (QL) F-test (using the functions glmQLFit and glmQLFTest). First, lowly expressed genes were filtered using filterByExpr function and then batch correction was performed using the additive model formulas in edgeR. Significantly differentially expressed genes were those having an FDR < 0.01 for BR, ET, JA and SA time series RNA-seq datasets and validation RNA-seq datasets (ABA, ET, JA, SA) or FDR < 0.05 with no batch effect correction for the time series ABA and SL/KAR datasets (Robinson et al., 2010).

For TF family enrichment analysis, the hypergeometric distribution was performed using phyper function in R. The distributions with p-value < 0.01 were considered significant. Known Arabidopsis TF information was obtained from PlantTFDB 5.0 (Jin et al., 2016). To estimate the significance of overlap between any two hormone treatments, a 2*2 table was generated as described in Nemhauser et al. (2006), and the Chi-square test (using chisq.test function in R) was performed based on the table. Clust analysis was performed according to Abu-Jamous and Kelly (2018). Heatmap analyses were performed by pheatmap with default parameters (https://github.com/raivokolde/pheatmap). The expression data 4 hours after hormone treatments were used for plotting the heatmaps because 2 hours data for ABA and ET treatments were not available.

### Hub target gene identification

Hub target genes were identified from networks of 17 hormone-relevant TFs built from ChIP-seq data generated by ourselves and two published studies (Song et al., 2016; Zander et al., 2020). All the ChIP-seq data used for constructing each hormone transcriptional network is described in detail in Supplementary Table 8 (ST. 8_1). Target genes in these networks were binned by the number of TFs that bind them. Target genes bound by more than 7 TFs were defined as hub target genes. The remaining genes were non-hub genes and were divided into the other two groups (group low, genes that are targeted by 1-3 TFs; group moderate, genes that are targeted by 4-6 TFs.).

For differential expression density plots, log2 fold changes in gene expression relative to 0 h were calculated. The expression data 2 hours after BR, JA, SA and SL/KAR treatment were used for consistency with ChIP-seq data, but 4 hours after ABA and ET treatments because 2 hours data were not available. The p-value was calculated by two-sample K-S test to indicate the distribution difference (p-value < 0.05).

### Signaling and dynamic regulatory events modeling

Regulatory networks were modeled using the SDREM framework (Gitter and Bar-Joseph, 2016). SDREM modeling uses as input condition specific time series transcriptomes and general information about hormone receptors identities TF-gene interactions and protein-protein interactions (PPIs) data.

For the time series transcriptomes, Log2 fold changes relative to 0 h for all expressed genes were calculated at 15 min, 30 min, 1 h, 2 h, 4 h, 8 h, 12 h, 24 h post-hormone treatment. TF-target gene interactions were collected from several sources. First, the TF-gene interactions were identified from ChIP-seq data for 17 TFs. The detailed information of which data was used for constructing SDREM models for each hormone is described in Supplementary Table 8 (ST. 8_1). Second, 406,832 TF-gene interactions for 296 TFs were included from published DAP-seq studies (O’Malley et al., 2016; Narsai et al., 2017; Zander et al., 2020). Third, confirmed and direct 4,378 interactions for 293 TFs were obtained from the Arabidopsis Gene Regulatory Information Server repository (Yilmaz et al., 2010).

PPIs were obtained from BioGRID and the combined phytohormone interactome network (Stark et al., 2006; Altmann et al., 2020). The PPI weight score was applied according to SDREM methods (Gitter and Bar-Joseph, 2016).

The identities of hormone receptors for each hormone are listed in Supplementary Data 2 (files: aba_source.txt, br_source.txt, et_source.txt, ja_source.txt, sa_source.txt, sl_kar_source.txt). The receptors for ABA were 14 PYR/PYL/RCARs (Ma et al., 2009; Park et al., 2009), for ET were ETR1, 2, ERS1, 2 and EIN4 (Bleecker et al., 1988; Chang et al., 1993; Hua et al., 1995; Hua et al., 1998; Sakai et al., 1998), for SA were NPR1, 2, 3 (Castelló et al., 2018; Ding et al., 2018), and for SL/KAR were AtD14 and KAI2 (Waters et al., 2012). The receptor for BR was BRI1 (Clouse et al., 1996; Wang et al., 2001), and for JA was COI1 (Xie et al., 1998).

When running SDREM the Minimum_Absolute_Log_Ratio_Expression parameter was used at default parameter 1, which retains only the genes whose largest absolute log2 fold changes across all time points is greater than 1. The maximum path length parameter was set to 5 and only binary splits were allowed in the regulatory paths. SDREM was run for 10 iterations for ABA, BR, JA, SA and SL/KAR models. We extended extra 2 iterations for ET SDREM model as the TFs and signaling proteins predicted in each iteration did not substantially converge across iterations when only running 10 iterations. The parameters used in modified DREM and SDREM modeling are given in Supplementary Data 1, 2. SDREM reconstructed signaling pathway results were visualized in the tool and by using Cytoscape v3.8.0. Intersection analysis was conducted using Intervene (Khan and Mathelier, 2017). The integrated transcriptional cross-regulation model was generated by overlaying the individual hormone transcriptional regulatory models.

### Functional enrichment analyses

Gene ontology (GO) enrichment analysis of predicted nodes in each hormone model was conducted using the compareCluster function in clusterProfiler with default parameters (Yu et al., 2012).

The functional grouped network of hub genes (group; hubs) and non-hub genes (groups; low and moderate) was performed using ClueGO v2.5.7 in Cytoscape v3.8.0 with ontologies were updated as GO_MolecularFunction-Custom-GOA-ACAP-ARAP_28.08.2020 (Bindea et al., 2009). Benjamini-Hochberg was used to correct the p-values for multiple testing. Functional groups with a p-value < 0.01 were considered statistically significantly enriched. The network specificity was set to ‘Global’. GO term fusion was selected to reduce terms redundancy. The kappa score was set as ≥ 0.4 to connect the terms in the network. All the settings above were kept same for the three groups.

ClueGO V2.5.7 was used for GO enrichment analysis of the 23 proteins shared by at least four hormone signaling pathways. The ontologies were updated as GO_MolecularFunction-Custom-GOA-ACAP-ARAP_28.08.2020; GO_CellularComponent-Custom-GOA-ACAP-ARAP_28.08.2020 and GO_BiologicalProcess-Custom-GOA-ACAP-ARAP_28.08.2020. Benjamini-Hochberg was selected to correct the p-values for multiple testing corrections. Functional groups with a p-value < 0.05 were considered statistically significantly enriched. The network specificity was set to ‘Global’ with minimum 5 genes/term. GO term fusion was selected to reduce term redundancy. The other settings were kept as default.

### Protein extraction and digestion

Peptides for quantitative protein abundance estimation were generated using a filter-aided sample preparation (FASP) method followed by on filter digestion as previously described with certain modifications (Song et al., 2020). Frozen plant tissue was ground to a fine powder (∼0.3 g per sample), then proteins were extracted by adding 5 volumes of protein extraction buffer containing 1X MS-SAFE protease and phosphatase inhibitor (Sigma MSSAFE) and 1mM PMSF. Proteins were precipitated using pre-chilled methanol containing 0.1M Ammonium acetate. Pellets were resuspended in 1mL of resuspension buffer (8M Urea, 50mM Tris pH 7.5 and 5mM Tris(2-carboxyethyl) phosphine hydrochloride (TCEP)) using a bath sonicator for 30 mins. Protein concentration was determined using BCA assay (ThermoFisher Scientific 23225). FASP was performed using Amicon Ultracel-30K centrifugal filters -4mL (Millipore UFC803008). Proteins were digested by adding bovine trypsin (Worthington-LS003750) and endoproteinase Lys-C (NEB-P8109S) at a ratio of (1:25) and (1:800) respectively for each sample and incubated at 37°C overnight. Digested peptides were desalted using sep-pak C18 Cartridge (Waters-WAT051910) conditioned as per manufacturer’s protocol. Peptides were eluted stepwise with 1mL of 20% Acetonitrile-water, 1mL of 40% Acetonitrile-water and 2ml of 80% Acetonitrile-water. Eluates were combined and peptide content was estimated using protein A280 on a NanoDrop 2000 (Thermo Scientific). Eluate was split into two fractions with the smaller fraction (∼5µg of peptide) reserved for total protein abundance and the larger fraction taken forward for phosphopeptide enrichment. Both fractions were dried using speedvac.

### Phosphopeptide enrichment

Phosphopeptides were enriched using the High SelectTM TiO_2_ phosphopeptide enrichment kit (A32993-ThermoFisher Scientific) Briefly, dried peptides were resuspended in 150µL of binding and equilibration buffer. The TiO_2_ zip-tips were conditioned using 20µL wash buffer followed by 20µL of binding and equilibration buffer and centrifuged at 3000 xg for 2min. Peptide solution was applied to the zip-tip and centrifuged at 1000 xg for 5 min. Flow-through was collected and reapplied onto the zip-tip. The zip-tips were washed with 20µL binding and equilibration buffer followed by 20µL of wash buffer, and this was repeated once more, followed by a final wash with 20µL of LC-MS grade water. Liquid was removed by centrifugation and excess liquid was blot dried on a clean lab tissue. The zip-tips were transferred into a fresh collection tube and phosphopeptides were eluted using 50µL of elution buffer twice. Eluates were dried in a speedvac.

### Liquid chromatography-tandem mass spectrometry (LC-MS/MS) analysis

Proteomic and phosphoproteomic samples were analyzed by LC-MS/MS using a Thermo Ultimate 3000 RSLCnano UHPLC system connected to a Thermo Orbitrap Eclipse Tribrid mass spectrometer (Thermo-Fisher Scientific, Waltham, MA, USA). Peptides were reconstituted in 0.1% trifluoroacetic acid (TFA) and 2% acetonitrile (ACN) and loaded onto a PepMap C18 5 µm 1 cm trapping cartridge (Thermo-Fisher Scientific, Waltham, MA, USA) at 12 µL/min for 6 min before switching the pre-column in line with the analytical column (nanoEase M/Z Peptide BEH C18 Column, 1.7 µm, 130 Å and 75 µm ID × 25 cm, Waters). The column compartment was held at 55°C for the entire analysis. Separation of peptides was performed at 250 nL/min using a linear ACN gradient of buffer A (0.1% formic acid, 2% ACN) and buffer B (0.1% formic acid, 80% ACN), starting at 14% buffer B to 35% over 90 min, then rising to 50% B over 15 min. Buffer B was ramped up to 95% in 5 min and the column was cleaned for 5 min at 95% B followed by a 3 min equilibration step at 1% B.

Mass spectra were collected in Data Dependent Acquisition (DDA) mode. MS1 spectra were collected in the Orbitrap while HCD MS2 spectra were collected in the ion trap. MS1 scan parameters: scan range of 375-1650 m/z, 120,000 resolution, max injection time of 50 ms, AGC target 4e5, HCD collision energy 30%. Easy-IC internal mass calibration was used. MS2 spectra were collected in the ion trap on rapid mode, AGC target of 1e4, max IT of 35 ms. The isolation window of the quadrupole for the precursor was 0.8. Dynamic exclusion parameters were set as follows: exclude isotope on, duration 60 s and using the peptide monoisotopic peak determination mode, charge states of 2-7 were included.

### Raw mass spectrometry data processing and differential expression analysis

Raw files obtained from mass spectrometry analysis were searched against the *Arabidopsis thaliana* protein sequence database (version TAIR10) obtained from the TAIR website, using Sequest HT through Proteome Discoverer (Version 2.4) (Thermo Scientific, Bremen, Germany). Precursor and fragment mass tolerance were set to 20 ppm and 0.5 Da, respectively. Carbamidomethylation of cysteine was set as fixed modification, while oxidation of methionine, acetylation of the protein N-terminus and phosphorylation at serine, threonine and tyrosine were set as dynamic modifications. A false discovery rate (FDR) threshold of 1% was used to filter peptide spectrum matches (PSMs). FDR was calculated using a concatenated target/decoy strategy in Percolator. For label-free quantification, precursor peaks were detected and quantified using the Minora Feature Detector and Precursor Ions Quantifier respectively. Data were normalized based on the total peptide amount using the normalization feature available in the Precursor Ions Quantifier node. A phosphoRS score threshold of ≥ 75% was used for phosphosite localization. Differentially expressed proteins and phospho-sites were identified using PoissonSeq with a p-value cut-off of 0.05 and fold change > 1.1. Sample loading normalization was performed before differential expression analysis.

### Identification of differentially alternatively spliced genes and isoform switch events

To detect differentially alternatively spliced genes, the union pipeline was used (Guo et al., 2020). Only expressed transcripts that had ≥ 1 counts per million (CPM) in one or more samples were retained. Read counts were normalized by the Trimmed Mean of M-values (TMM) method using edgeR (Robinson et al., 2010). Batch effects were estimated and removed using RUVSeq R package with the remove unwanted variations (RUVs) approach (Risso et al., 2014). Then the voom-weight function in limma and DiffSplice functions were used for differentially alternatively spliced analysis (Ritchie et al., 2015).

Significantly differentially alternatively spliced genes were determined by using the following criteria. Firstly, at least one of the transcripts differed significantly in log2 fold changes from the corresponding gene with an adjusted p-value of < 0.05, and secondly at least one of the transcripts of the gene exhibited Δ percent spliced (Δ PS) ≥ 0.1. The PS value was estimated as the ratio of a transcript’s average abundance divided by the average of its corresponding gene abundance. The SF-RBPs list was obtained from Calixto et al. (2018).

For detection of alternatively spliced isoform-switch events, the TSIS R package was used with time-series transcriptome data as described previously (Zander et al., 2020). Transcripts with average TPM across all time points > 1 were included in the TSIS analysis. The mean expression approach was used to search for interaction points. Statistically significant switch events were identified using the following filtering parameters: (1) probability cut-off value of > 0.5; (2) differences cut-off value of > 1; (3) *p* cut-off value of < 0.05; (4) minimum time in interval of > 1. The protein-coding transcripts information was obtained from Zhang et al. (2017).

### Statistics

For estimating TF family enrichment significance, the hypergeometric test was performed using phyper function in R. The Chi-square test was used to estimate the significance of overlap between any two hormone treatments. The two-sample K-S test was used for testing the distribution difference of expression density plots using ks.test in R. To test differentially expressed genes in RNA-seq, the significance was calculated from a quasi-likelihood (QL) F-test and corrected with Benjamini-Hochberg correction for multiple testing using edgeR 3.28. For proteomics experiments, we used the p cutoffs generated from the statistical tests based on reference (Clark et al., 2021).

## Data availability

ChIP-seq data in WT (Col-0) and transgenic seedlings, and validation RNA-seq data for ABA, ET, JA and SA in WT and *mpk6* mutant seedlings generated in this study can be downloaded from the Gene Expression Omnibus repository (GEO, https://www.ncbi.nlm.nih.gov/geo/) with accession number GSE220957 and reviewer token qvarqgwwdnoljob. RNA-seq data for time series BR, SA, SL/KAR can be downloaded from GEO with accession number GSE182617 and reviewer token qrunyeymtpurngt. The mass spectrometry proteomics data have been deposited to the ProteomeXchange Consortium *via* the PRIDE partner repository with the dataset identifier PXD039958. Reviewer username is: and the password is: AmCP2SU6. ET RNA-seq raw reads were downloaded from Sequence Read Archive (SRA, https://www.ncbi.nlm.nih.gov/sra) with accession number SRA063695. ABA RNA-seq raw reads and ChIP-seq raw reads for ABF1, 3, 4 were downloaded from GEO with accession number GSE80568. JA RNA-seq raw reads were downloaded from GEO with accession number GSE133408. The PPIs with applied scores, the TF-target interaction inputs, the parameters and the output models for recreating models for each hormone in this study can be found in Supplementary Data 1, 2 in the Source_data folder. All the code used for this study can be found in GitHub: https://github.com/LynnYin7911/hormone-network.

## Supporting information

Source Data for SDREM model reconstruction

## Acknowledgements

MGL was supported by an EU Marie Curie FP7 International Outgoing Fellowship (252475). This work was supported by grants from the National Science Foundation (NSF) (MCB-1024999 to JRE), the National Institutes of Health (R01GM120316), the Division of Chemical Sciences, Geosciences, and Biosciences, the Office of Basic Energy Sciences of the US Department of Energy (DE-FG02-04ER15517), and the Gordon and Betty Moore Foundation (GBMF3034). JRE is an Investigator of the Howard Hughes Medical Institute. MGL’s lab is funded by the Australian Research Council (ARC) Discovery Program grant DP220102840. We thank Jeff A. Long for providing recombineering reagents and protocols. We thank the following undergraduates, technicians and staff scientists who contributed technical assistance to the project: R Carlos-Serrano, L. Tames, J. Park, O. Romero, R. Luong, W. Ho, Y. Koga, S. Hazelton, H. Chen and M. Urich. We thank the La Trobe Proteomics platform, Keshava K Datta and Rohan Lowe for processing (phospho)proteomic samples. We thank Aaron Wise for help establishing SDREM analyses and Anthony Gitter for ongoing advice on refining SDREM analyses.

## Author contributions

MGL, JRE and LY designed the study. JAL provided novel reagents and methods. MGL, MZ, MX, LS, JPSG, EH, RCS and BJJ generated the transgenic constructs and carried out the time series RNA-seq and ChIP-seq lab experiments. SCH, SJ, AW and ZB-J provided new analytical tools and analyzed data. SN, BKS, TB, JW, NMC and LY planned, conducted and analyzed the *mpk6* experiments. LY analyzed and integrated all data, interpreted results and prepared figures. LY and MGL drafted the manuscript. All authors read and approved the final manuscript.

## Supplementary Table Legends

**Supplementary Table 1.** Differentially expressed genes for each hormone response time series, determined by RNA-seq.

**Supplementary Table 2.** Overview of quality metrics of generated ChIP-seq datasets by this study.

**Supplementary Table 3.** Target genes and binding sites of 17 TFs under air or hormone treatments, determined by ChIP-seq.

**Supplementary Table 4.** The regulatory network determined by modified DREM models for all hormones.

**Supplementary Table 5.** Overview of six hormone signaling pathways reconstructed by SDREM modeling.

**Supplementary Table 6.** The predicted active TFs and their family distributions for each hormone.

**Supplementary Table 7.** List of number of differentially expressed genes shared between any two hormone response transcriptomes and overview of the differentially expressed TFs and their family distributions in each hormone dataset.

**Supplementary Table 8.** Overview of hub target genes and non-hub genes bound by 17 hormone TFs in hormone transcriptional networks.

**Supplementary Table 9.** Differentially expressed proteins detected in proteomics analyses.

**Supplementary Table 10.** Differentially abundant phosphopeptides detected in phosphoproteomics analyses.

**Supplementary Table 11.** The shared and unique differentially expressed proteins and phosphopeptides between two genotypes.

**Supplementary Table 12.** Differentially expressed genes for 1 hour hormone response (ABA, ET, JA, SA) identified by RNA-seq.

**Supplementary Table 13.** Overview of the differentially alternatively spliced genes for each hormone response time series analyzed, determined by transcript-level time series RNA-seq.

**Supplementary Table 14.** Overview of the isoform switch events in the time series RNA-seq for each hormone analyzed.

**Supplementary Table 15.** Overview of the first appear differentially expressed genes and differentially alternatively spliced genes at each time point.

## Supplementary Data Legends

**Supplementary Data 1.** The inputs, parameters and output models for recreating the regulatory networks for each hormone.

**Supplementary Data 2.** The inputs and parameters for recreating the signaling pathways for each hormone.

**Extended Data Figure 1.**
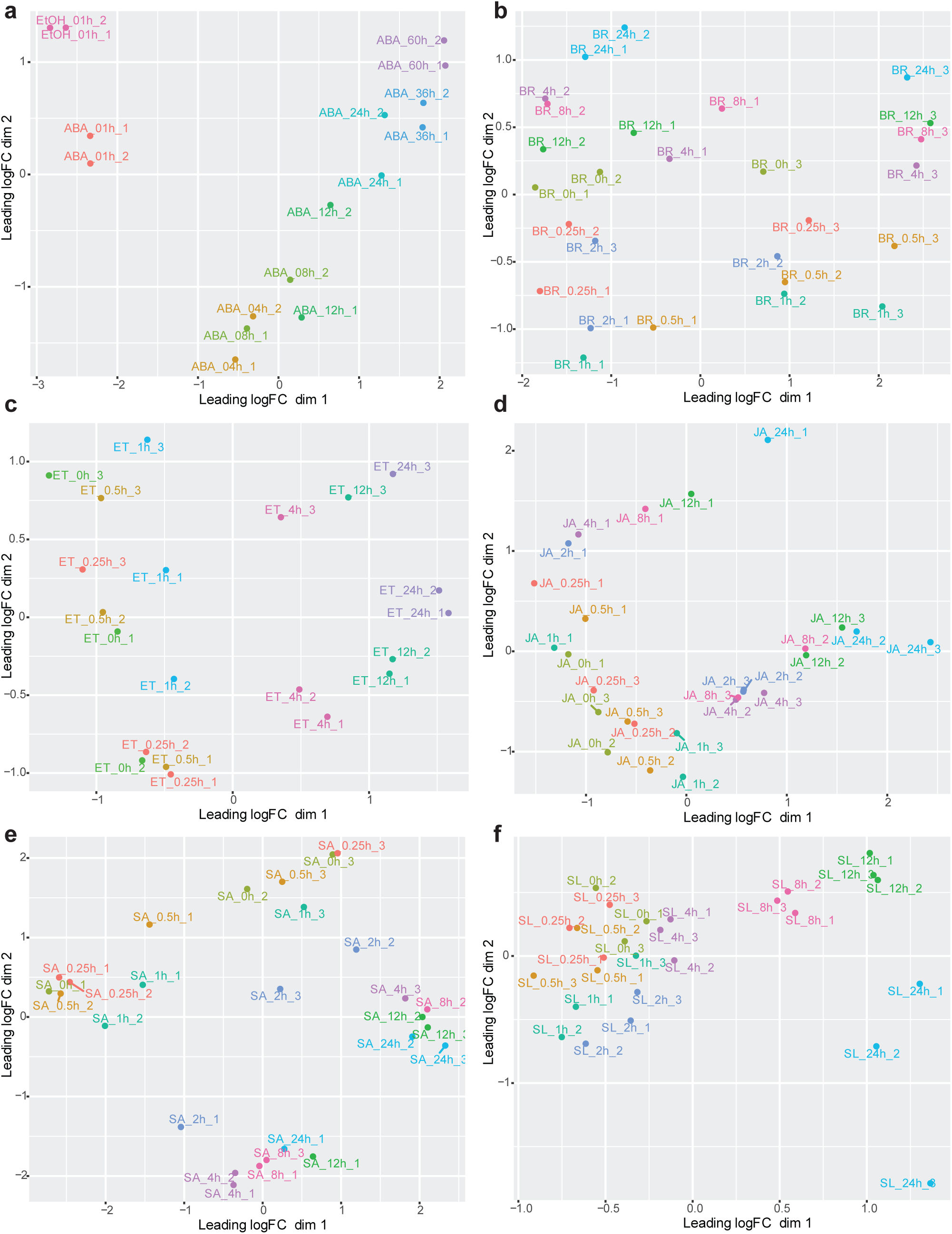
Overview of quality metrics of RNA-seq data. **a-f**, Multidimensional scaling (MDS) plots of replicate samples of the ABA, BR, ET, JA, SA and SL/KAR (labelled here as SL) treatment RNA-seq time-series in WT. BR, ET, JA, SA and SL/KAR treatment time series consist of three independent samples (n = 3) for each time point. ABA treatment time series consist of two independent samples (n = 2) for each time point. The ethanol treated 1 hour sample (EtOH_01h) was used as the mock treated samples for ABA treated samples when performing the differentially expressed gene analysis, because no 0 hour samples were collected during the experiment.

**Extended Data Figure 2.**
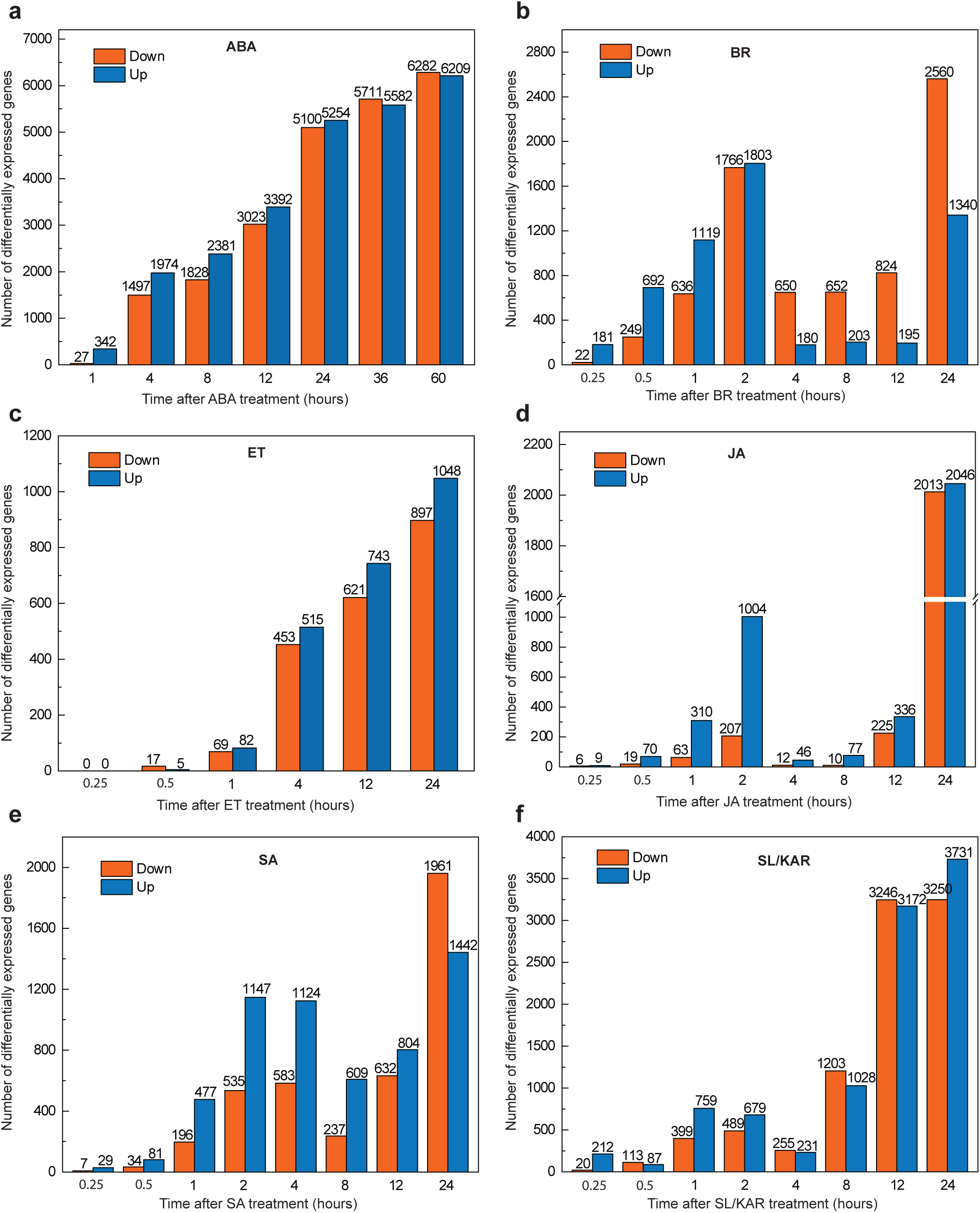
Time-series transcriptome analysis. **a-f**, Plots show the numbers of significantly differentially expressed genes (edgeR; FDR < 0.01 for BR, ET, JA and SA datasets; FDR < 0.05 for ABA and SL/KAR datasets) relative to 0 h upon hormone treatment. The x-axis represents time after hormone treatment (hours). The y-axis represents the numbers of significantly down-regulated and up-regulated genes, which are represented by orange and blue bars respectively.

**Extended Data Figure 3.**
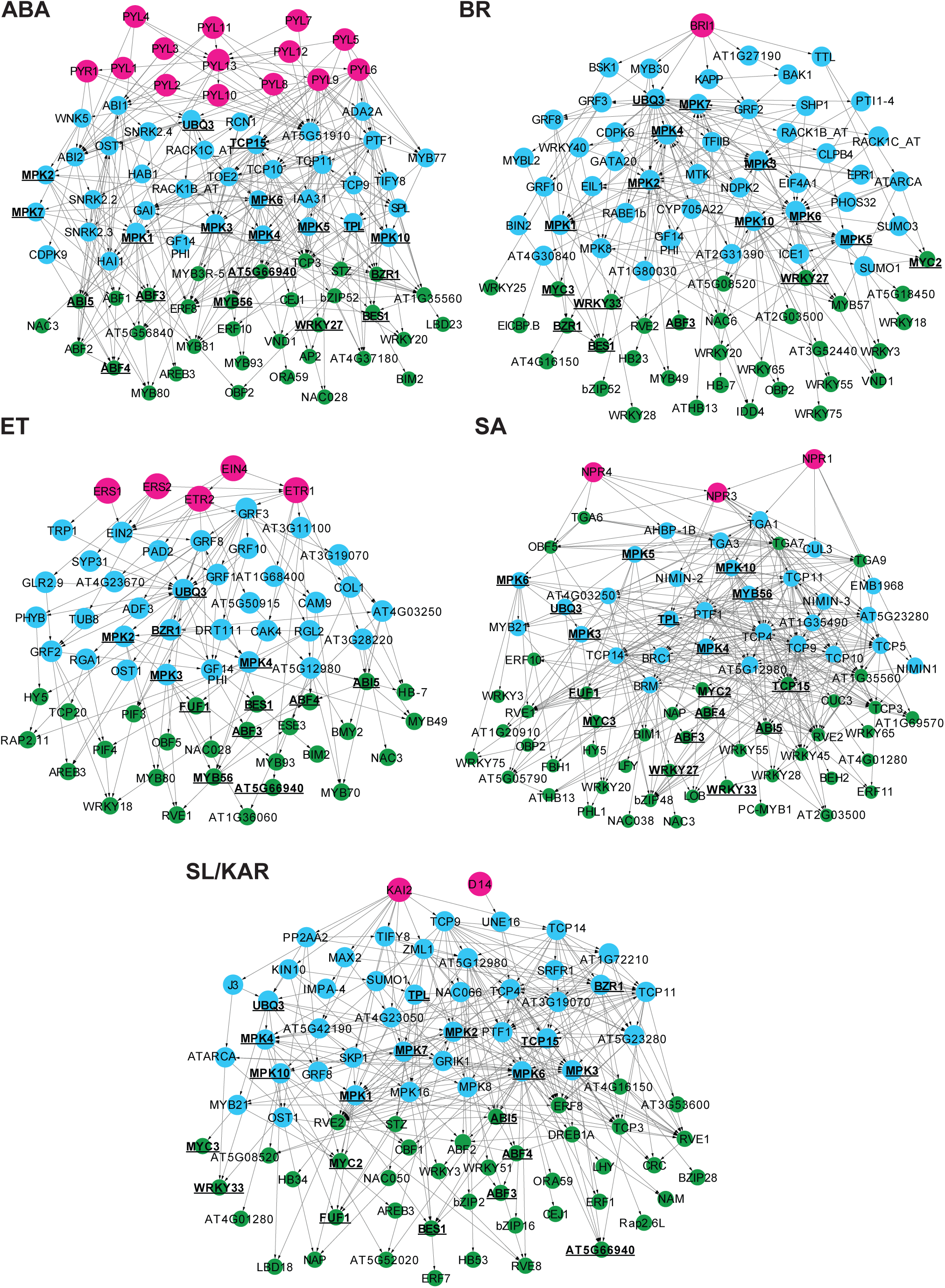
The reconstructed transcriptional regulatory models for ABA, BR, ET, SA and SL/KAR. The hormone receptor(s), intermediate proteins and active TFs are represented by magenta, blue and green nodes respectively. The proteins shared by at least 4 hormone pathways are in black bold text and have underlined names.

**Extended Data Figure 4.**
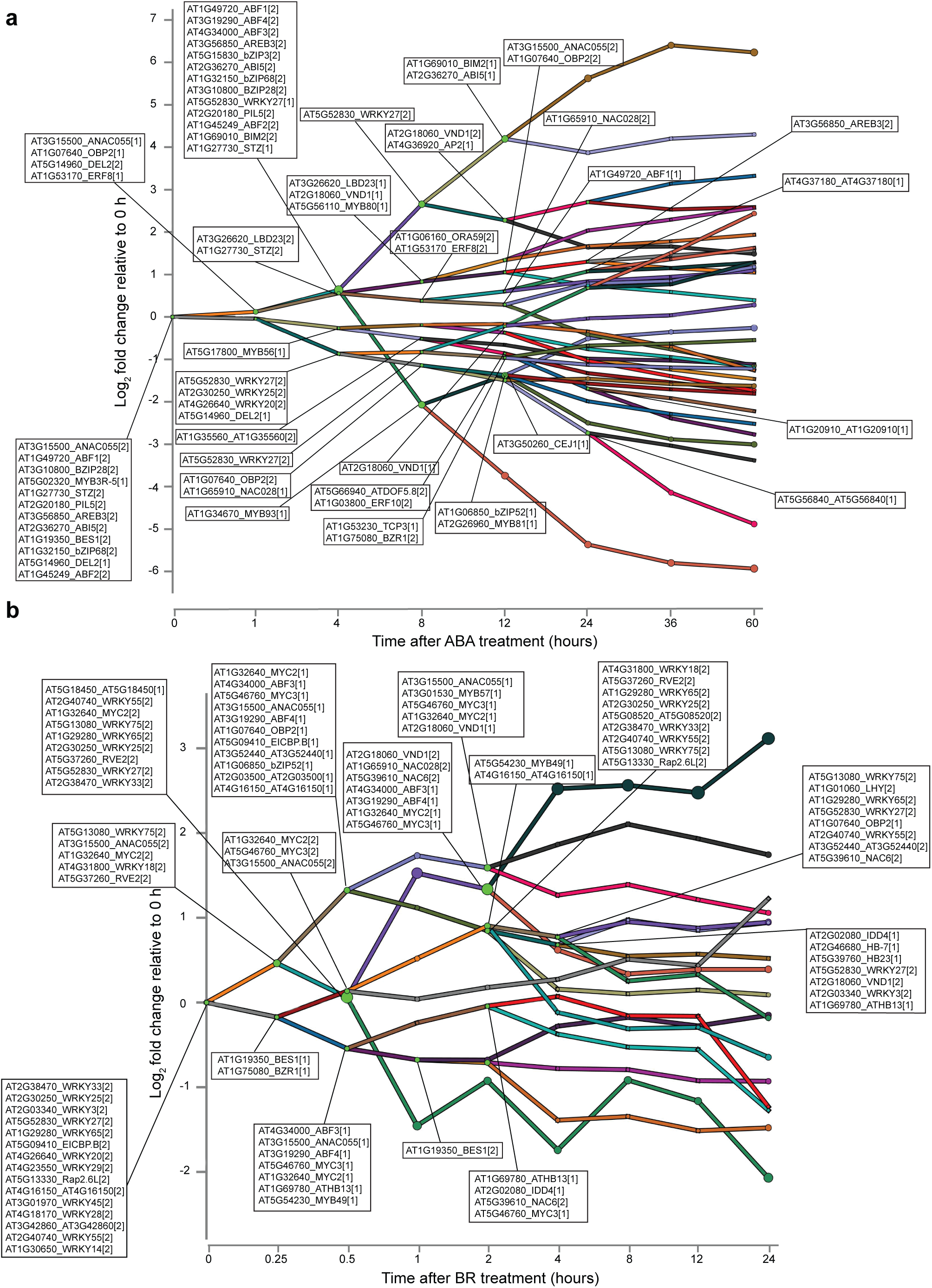
The transcriptional regulatory network component of the ABA (a) and BR (b) models. The networks display all predicted active TFs at each branch point (node) and the bars indicate co-expressed and co-regulated genes for each hormone. [1] indicates the TF primarily controls the lower path out of the split and [2] is for the higher path. The y-axis is the log2 fold change in expression relative to expression at 0 h.

**Extended Data Figure 5.**
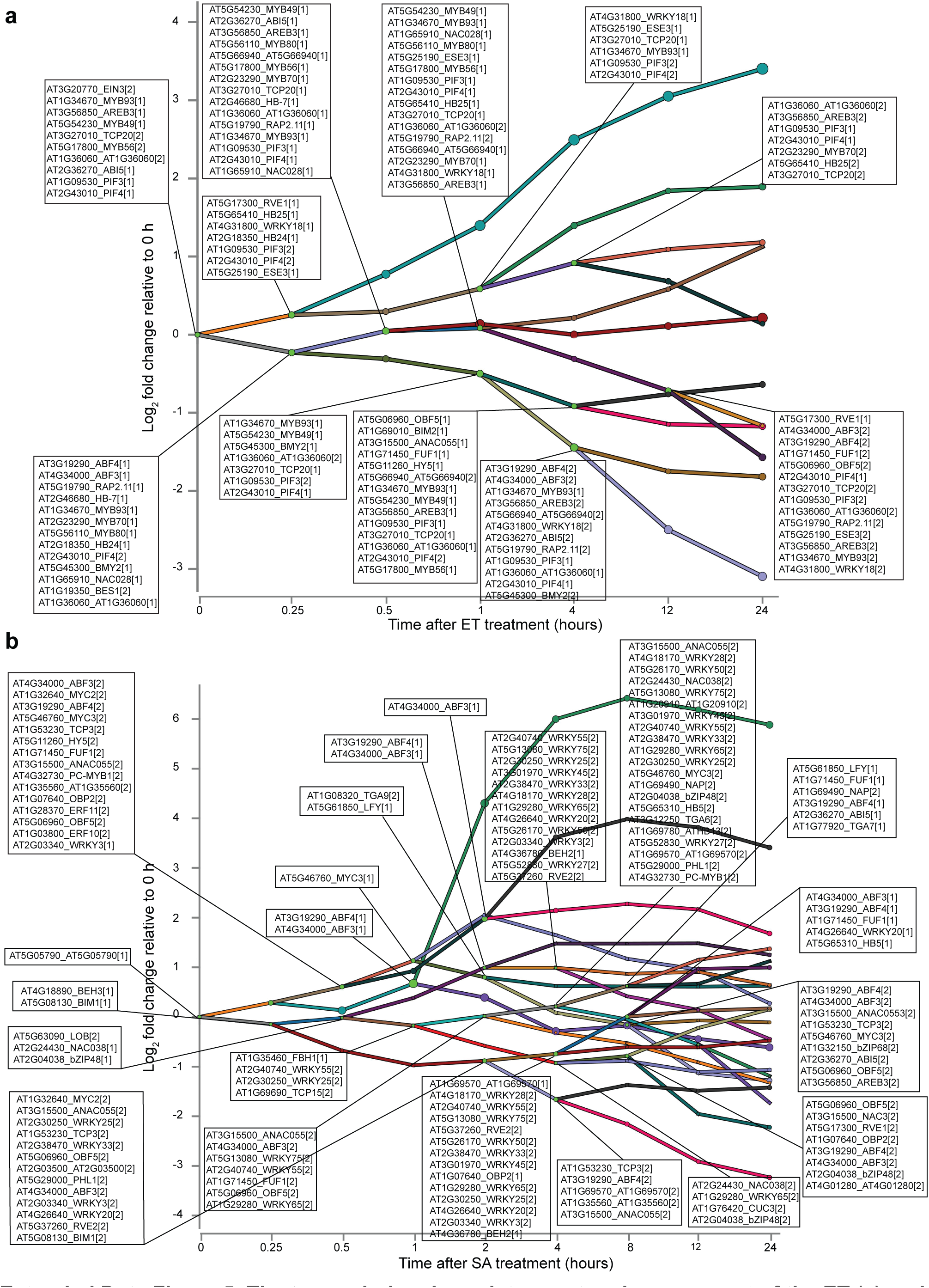
The transcriptional regulatory network component of the ET (a) and SA (b) models. The networks display all predicted active TFs at each branch point (node) and the bars indicate co-expressed and co-regulated genes for each hormone. [1] indicates the TF primarily controls the lower path out of the split and [2] is for the higher path. The y-axis is the log2 fold change in expression relative to expression at 0 h.

**Extended Data Figure 6.**
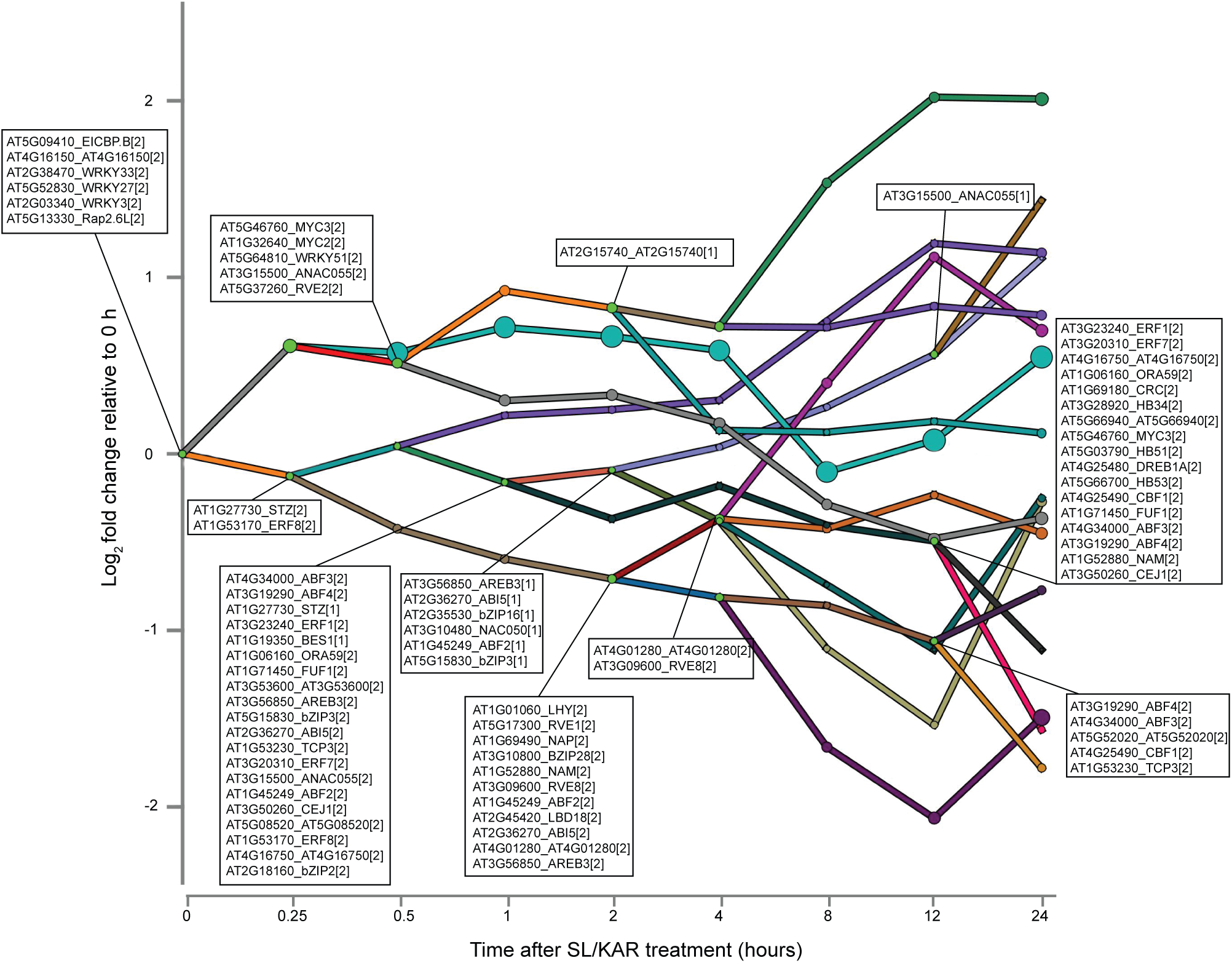
The transcriptional regulatory network component of the SL/KAR model. The network display all predicted active TFs at each branch point (node) and the bars indicate co-expressed and co-regulated genes. [1] indicates the TF primarily controls the lower path out of the split and [2] is for the higher path. The y-axis is the log2 fold change in expression relative to expression at 0 h.

**Extended Data Figure 7.**
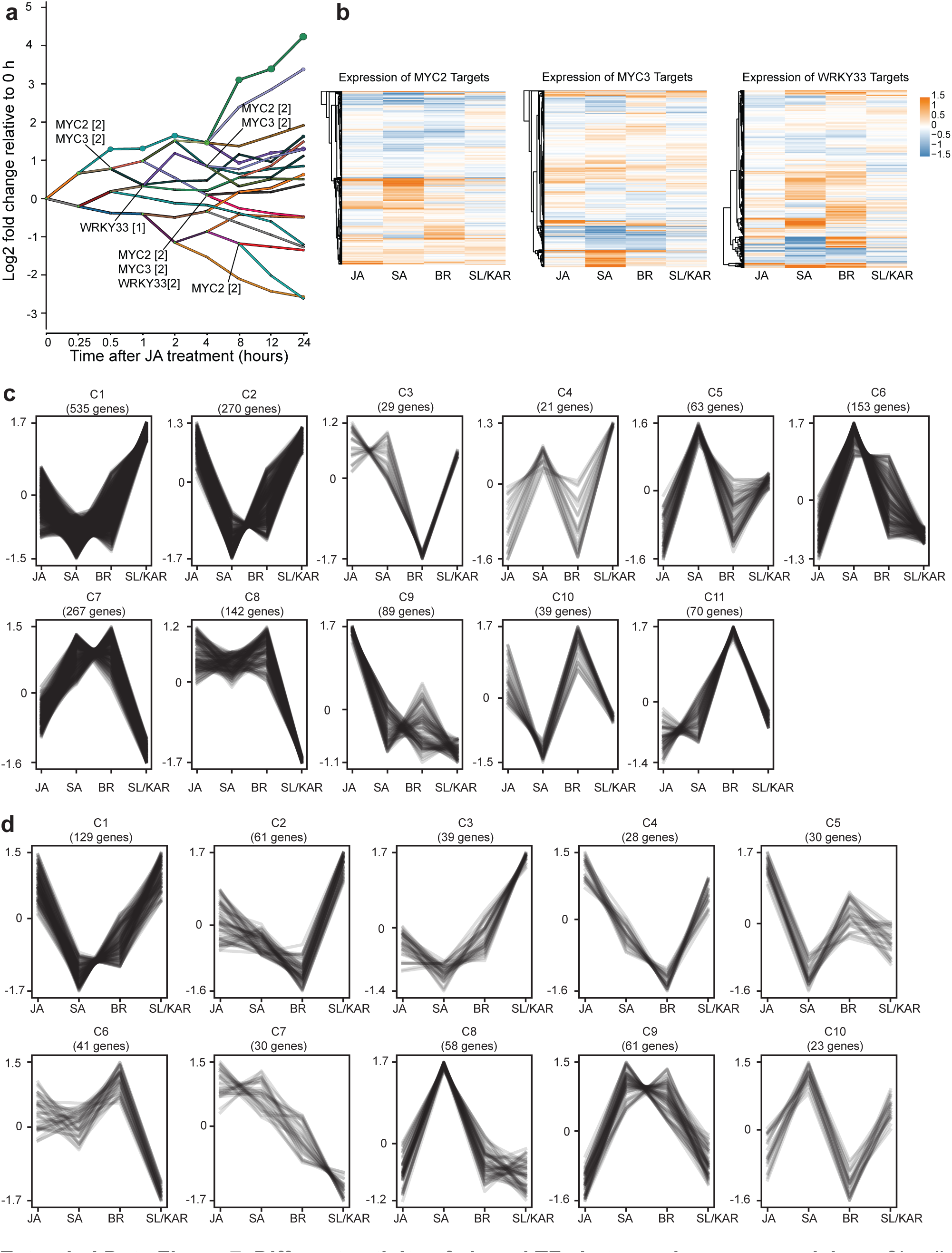
Different activity of shared TFs between hormone models. **a**, Simplified transcriptional regulatory network component for JA response highlighting the association of MYC2, MYC3 and WRKY33 with up and down regulated genes. **b**, Heatmap of expression of targets for MYC2, MYC3 and WRKY33 during JA, SA, BR and SL/KAR hormone responses. Hormone models are indicated in the x-axis, expression is given as log2 fold change relative to 0 h. **c, d**, K-means clustering of expression of MYC3 (c) and WRKY33 (d) target genes during JA, SA, BR and SL/KAR hormone responses. Expression is given as normalized transcripts per million (TPM).

**Extended Data Figure 8.**
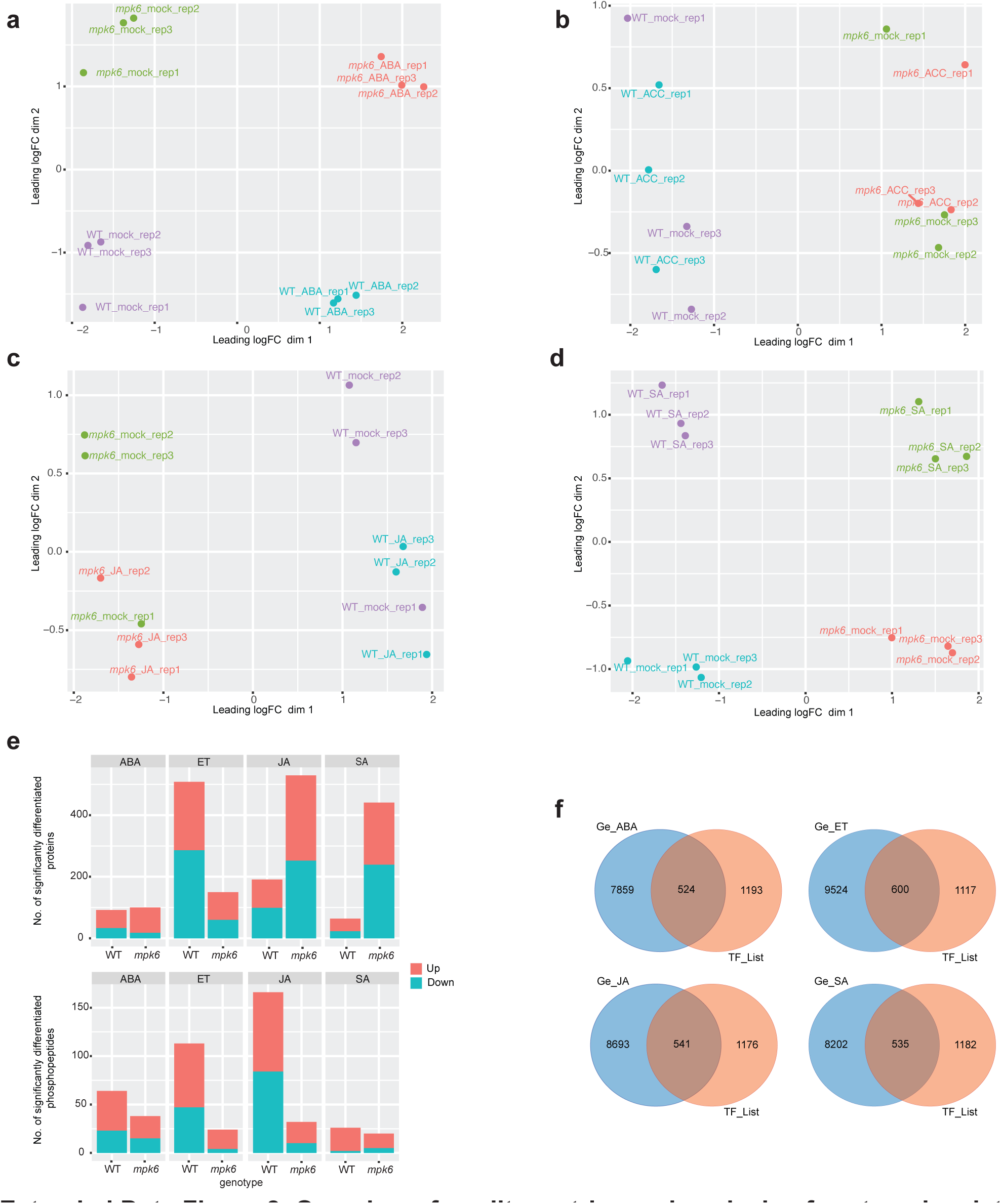
Overview of quality metrics and analysis of proteomics data. **a-d**, Multidimensional scaling (MDS) plots of replicate samples of the ABA, ET, JA, SA treatment for 1 h RNA-seq in WT and *mpk6* seedlings. All hormone treatments consist of three independent samples (n = 3). **e**, Total number of significantly differentially abundant proteins and phosphopeptides detected in comparisons between hormone-treated (ABA, ET, JA, SA; 1 h) WT and *mpk6* seedlings and mock controls (p-value < 0.05 & fold change > 1.1). Three independent experiments (with or without 1 h of hormone treatment; n = 3) were conducted for WT and *mpk6* seedlings. **f**, The number of TFs presents amongst the differentially expressed genes detected in comparisons between hormone-treated (ABA, ET, JA, SA; 1 h) *mpk6* and hormone-treated WT seedlings. (Ge_ABA: *mpk6* ABA versus WT ABA; Ge_ET: *mpk6* ET versus WT ET; Ge_JA: *mpk6* JA versus WT JA; Ge_SA: *mpk6* SA versus WT SA; TF_List: Known Arabidopsis TFs which were obtained from PlantTFDB 5.0).

**Extended Data Figure 9.**
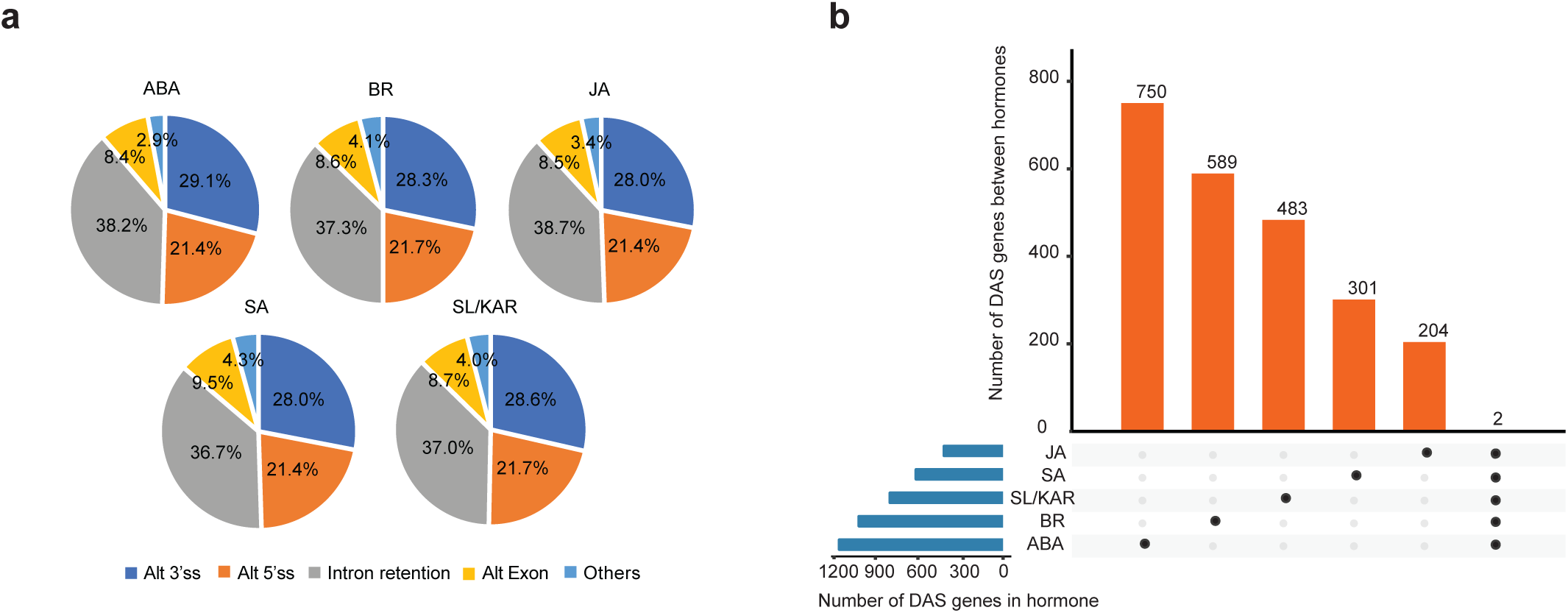
The number and alternative splicing types of the differentially alternative spliced genes in response to hormone. **a**, The major alternative splicing types of DAS genes in each hormone. Alt 3’ss: Alternative 3’ (A3) splice sites, Alt 5’ss: Alternative 5’ (A5) splice sites, Intron retention, Alt Exon: Skipping exon (SE) and Mutually Exclusive (MX) exons, Others: Alternative First (AF) and Last (AL) exons. **b**, The number of differentially alternative spliced (DAS) genes unique to and shared between all five hormones analyzed.

