## Supplementary material for "Transcription Factor Dynamics in Cross-Regulation of Plant Hormone Signaling Pathways": Source Data for SDREM model reconstruction: README.txt

1.The gene expression file (one input for running SDREM)
aba_DEG_matrix.txt
br_DEG_matrix.txt
et_DEG_matrix.txt
ja_DEG_matrix.txt
sa_DEG_matrix.txt
sl_kar_DEG_matrix.txt

2. The separate DREM properties file that used for running DREM software(as described in http://sb.cs.cmu.edu/drem/DREMmanual.pdf) except the TF-gene_Interactions_File will be generated dynamically and Active_TF_influence is a new SDREM parameter
aba_modified_DRME_parameter.txt
br_modified_DREM_parameters.txt
et_modified_DREM_parameters.txt
ja_modified_DREM_parameters.txt
sa_modified_DREM_parameters.txt
sl_kar_modified_DREM_parameters.txt


3.The DREM model file for iteration N (the final iteration of SDREM modeling), which can be loaded by drem.jar
aba_outmodel10.txt
br_outmodel10.txt
et_outmodel12.txt
ja_outmodel10.txt
sa_outmodel10.txt
sl_kar_outmodel10.txt

4.The new TF-gene binding file that will be used as input at next iteration. So here are the final TF-gene binding file in the SDREM modeling
aba_TFactive9.txt
br_TFactive9.txt
et_TFactive11.txt
ja_TFactive9.txt
sa_TFactive9.txt
sl_kar_TFactive9.txt
