## Supplementary material for "Transcription Factor Dynamics in Cross-Regulation of Plant Hormone Signaling Pathways": Source Data for SDREM model reconstruction: README.txt

1. The identities of hormone receptors
aba_source.txt: 14 PYR/PYL/RCARs receptors in abscisic acid (ABA) signalling pathway
br_source.txt: BRI1 receptor in brassinosteroid(BR) signalling pathway
et_source.txt: Five receptors (ETR1,2, ERS1,2, EIN4) in ethylene(ET) signalling pathway
ja_source.txt: COI1 receptor in jasmonic acid(JA) signalling pathway
sa_source.txt: Three NPR receptors (NPR1,2,3) in salicylic acid signalling pathway
sl_kar_source.txt: The receptor AtD14 in strigolactone(SL) signalling pathway and the receptor KAI2 in karrikin(KAR) signalling pathway

2.The protein-protein interaction file (one input for running SDREM)
protein_ppi.txt

3.The integrated network by overlaying six hormones's signalling network
union_network.cys: the file can be imported into Cytoscape for visualizing the final network

4.SDREM properties files for reconstructing signalling pathways for each hormone.
aba_sdremEgfPriors.props
br_sdremEgfPriors.props
et_sdremEgfPriors.props
ja_sdremEgfPriors.props
sa_sdremEgfPriors.props
sl_kar_sdremEgfPriors.props
